# Targeted photothrombotic stroke leads to disruptions in neurovascular coupling

**DOI:** 10.1101/2022.01.22.477341

**Authors:** Smrithi Sunil, John Jiang, Shashwat Shah, Sreekanth Kura, Kivilcim Kilic, Sefik Evren Erdener, Cenk Ayata, Anna Devor, David A. Boas

## Abstract

Functional neuroimaging, which measures hemodynamic responses to brain activity, has great potential for monitoring stroke patients. However, the neurophysiological interpretations of these hemodynamic signals remain a challenge as the stroke is likely to alter both neural activity and neurovascular coupling. To address this challenge, we simultaneously captured neural activity, through fluorescence calcium imaging, and hemodynamics, through intrinsic optical signal imaging, during longitudinal stroke recovery. We found that photothrombotic stroke to somatosensory forelimb region altered neurovascular coupling in the acute phase (2 days and 1 week post-stroke) within the affected forelimb and peri-infarct regions. Neurovascular coupling was reestablished in the chronic phase (4 weeks post-stroke), and acute recovery of neurovascular coupling predicted sensorimotor function. Stroke also resulted in increases in the power of global brain oscillations, which showed distinct patterns between calcium and hemodynamics. Increased calcium excitability in the contralesional hemisphere was associated with increased intrahemispheric connectivity. Additionally, acute increases in hemodynamic oscillations were associated with improved sensorimotor outcomes.

**Teaser:** Acute ischemic stroke leads to neurovascular uncoupling and the extent of early recoupling predicts sensorimotor recovery.

## Introduction

An ischemic stroke occurs due to interruption of blood flow caused by thrombosis or embolism of a blood vessel, which leads to a reduction or complete loss of blood supply to downstream areas. Loss of blood supply causes a starved oxygen environment and leads to cellular damage within minutes and ultimately to sensorimotor and cognitive impairments^1,2^. A majority of stroke patients survive the incident, however, most survivors are compromised in work capacity, the extent of which varies across patients from mild to severe impairments^3^. Some spontaneous recovery is typically seen in most patients in the months following injury and most post-stroke recovery currently relies heavily on rehabilitative treatments^4–6^.

Functional neuroimaging methods, such as functional magnetic resonance imaging (fMRI) and functional near-infrared spectroscopy (fNIRS), which measure the hemodynamic response to brain activity, have the potential for being valuable tools for monitoring and managing the recovery and treatment of stroke patients both in the acute and chronic phases of stroke recovery^7–9^. However, the hemodynamic responses post-stroke are almost always altered relative to those seen in healthy individuals. Blood-oxygen-level-dependent fMRI (BOLD-fMRI) studies have revealed that task-related cortical responses following stroke undergo pronounced alterations in amplitude and spatial extent of the BOLD signal in both the ipsilesional and the contralesional hemispheres^9–11^. Additionally, studies assessing resting-state functional connectivity obtained with MRI (fc-MRI) have shown that inter-hemispheric connections are altered in the early acute phase of stroke in humans^12,13^. Whether these hemodynamic response alterations reflect the underlying differences in neural function or simply a result of injury to the vasculature is still under active investigation. In other words, we do not know the effect of stroke on neurovascular coupling and thus are limited in our ability to use these valuable neuroimaging tools to study functional recovery in stroke survivors.

Neurovascular coupling (NVC) has been studied extensively in healthy subjects and there is a large body of evidence suggesting that neural activity is closely related to cerebral blood flow (CBF) and oxygen metabolism^14,15^. This tight coupling between neural activity and hemodynamics forms the basis of modern neuroimaging techniques that use the cerebrovascular changes caused by neural activation to map changes in function in the behaving human brain^16^. While NVC is maintained in the healthy brain, brain pathologies such as traumatic brain injury, Alzheimer’s disease, and stroke may lead to disruptions in the interactions between neural activity and CBF, leading to neurovascular uncoupling, thereby confounding interpretations of neuroimaging results^17,18^. Additionally, the effect of stroke on NVC has received limited attention and sometimes led to conflicting results^9,19,20^. Thus, there is a need for preclinical stroke models to evaluate the functional aspects of neurovascular recovery and to use these findings to improve the interpretations of human neuroimaging studies.

Preclinical animal models of stroke have been used extensively over the last few decades to understand the mechanisms involved in stroke recovery from molecular and cellular changes to large scale functional network reorganizations^21–23^. On a mesoscopic level, studies performing *in vivo* calcium fluorescence imaging of neural activity have shown activation reorganization and functional remapping of the affected brain regions in the peri-infarct zone longitudinally, bearing on physiological processes underlying the evolution of stroke in humans^24–27^. Additionally, intrinsic optical signal imaging (IOSI) has been used to assess global changes using resting state functional connectivity analysis and also local changes in response to functional activation^28,29^. To improve interpretations of human functional neuroimaging studies and to understand the underlying physiology that gives rise to the observed hemodynamic signals we need to obtain simultaneous measures of neural and hemodynamic parameters post-stroke. Moreover, these measures need to be obtained on a mesoscopic scale to understand both the local and global changes that result due to stroke, as well as cover the entire longitudinal recovery phase to capture both acute and chronic time points.

Prior work on functional recovery following ischemic stroke has focused either just on neural or hemodynamic activity changes or just the acute or chronic phase and to the best of our knowledge these measures have not yet been integrated to study neurovascular coupling during stroke recovery^24,28–31^. In this paper, we study the relationships between neural and hemodynamic activity in the affected and unaffected hemispheres during longitudinal stroke recovery. We have previously established an optimized stroke model that more closely mimicked the physiology of a human stroke by inducing a stroke in an awake animal, occluding a single arteriole, and eliciting a distinct core and peri-infarct region. Here, we show that our optimized stroke model together with wide-field neural calcium and hemodynamic imaging can be used to monitor neurovascular coupling longitudinally. Our results suggest that acute stroke leads to neurovascular uncoupling as assessed through activity correlations and the hemodynamic response function. This uncoupling was capable of spontaneous re-coupling, which depended on the extent of initial acute uncoupling. Furthermore, the extent of neurovascular re-coupling was associated with improved sensorimotor outcomes.

## Results

### Wide-field fluorescence and intrinsic optical signal imaging can simultaneously follow changes in neural calcium and hemodynamic activity after stroke

Neurovascular coupling has been studied extensively in healthy subjects in both humans and animal models. In rodents, wide-field fluorescence calcium and intrinsic optical hemodynamic signals have been imaged simultaneously to investigate the baseline relationships between neural activity and blood flow^32,33^. Imaging calcium dynamics using GCaMP has been used extensively over the last decade as a correlate and reliable metric of neural activity^34,35^. Here, we first implemented these techniques to show that wide-field optical imaging can be used to investigate the differential effects of stroke on neural calcium dynamics and cerebral blood volume assessed with changes in the concentration of oxy and deoxy hemoglobin (HbO and HbR respectively). Fig 1a shows a simplified schematic of the imaging system and the experimental timeline. We first assessed alterations to evoked responses during sensory stimulation after stroke. Sensory stimulation using air-puff to the forelimb was performed in a block design paradigm (Fig 1b) and included 5 sec of baseline, followed by 5 sec of 3 Hz stimulation, followed by 20 sec of rest before the next trial. Each trial was repeated 20 times in one session. Fig 1b shows an example of the spatial maps and time-course of stimulus induced response in a healthy mouse. The raw GCaMP fluorescence signal was corrected for hemodynamic crosstalk using a modified attenuation estimation method prior to analysis^32,36^ (Fig 1b, Supplementary Fig 1). Unilateral photothrombotic stroke to the forelimb somatosensory cortex of the right hemisphere led to a significant suppression of the evoked calcium and hemodynamic responses to air-puff stimulation of the contralateral (affected) forelimb within the affected hemisphere, while the responses in the unaffected hemisphere were preserved (Fig 1c,d). The largest suppression of the response occurred 2 days post-stroke with a slow return of the response by 4 weeks, albeit still suppressed compared to pre-stroke. At day 2 after stroke, GCaMP showed a 70% reduction in the response. At the same time, total hemoglobin (HbT) and HbO showed a 100% and 45% reduction in response, respectively, compared to pre-stroke baseline (Supplementary Fig 2a). By 4 weeks after stroke the responses within the affected hemisphere had returned to 50% of the pre-stroke value. Evoked responses to forelimb air-puff stimulation of the unaffected forelimb did not exhibit significant alterations in the contralateral (unaffected) hemisphere, however, the affected hemisphere showed suppressed responses (Fig 1e,f). Once suppressed, the affected hemisphere did not recover either GCaMP or hemodynamic responses to ipsilateral (unaffected) forelimb stimulation even at 4 weeks (Supplementary Fig 2b). Spatiotemporal maps of responses during baseline, stimulation, and recovery at each time point are shown in Supplementary Fig 3 for the same mouse shown in Fig 1c,e. Knowing the co-evolution of neural and hemodynamic responses can aid in better interpretations of the alterations observed in the hemodynamic fMRI signals after stroke. The results here show that wide-field fluorescence and intrinsic optical signal imaging following photothrombotic stroke are sensitive measures that allow the longitudinal monitoring of these neural and hemodynamic signals.

**Figure 1:**
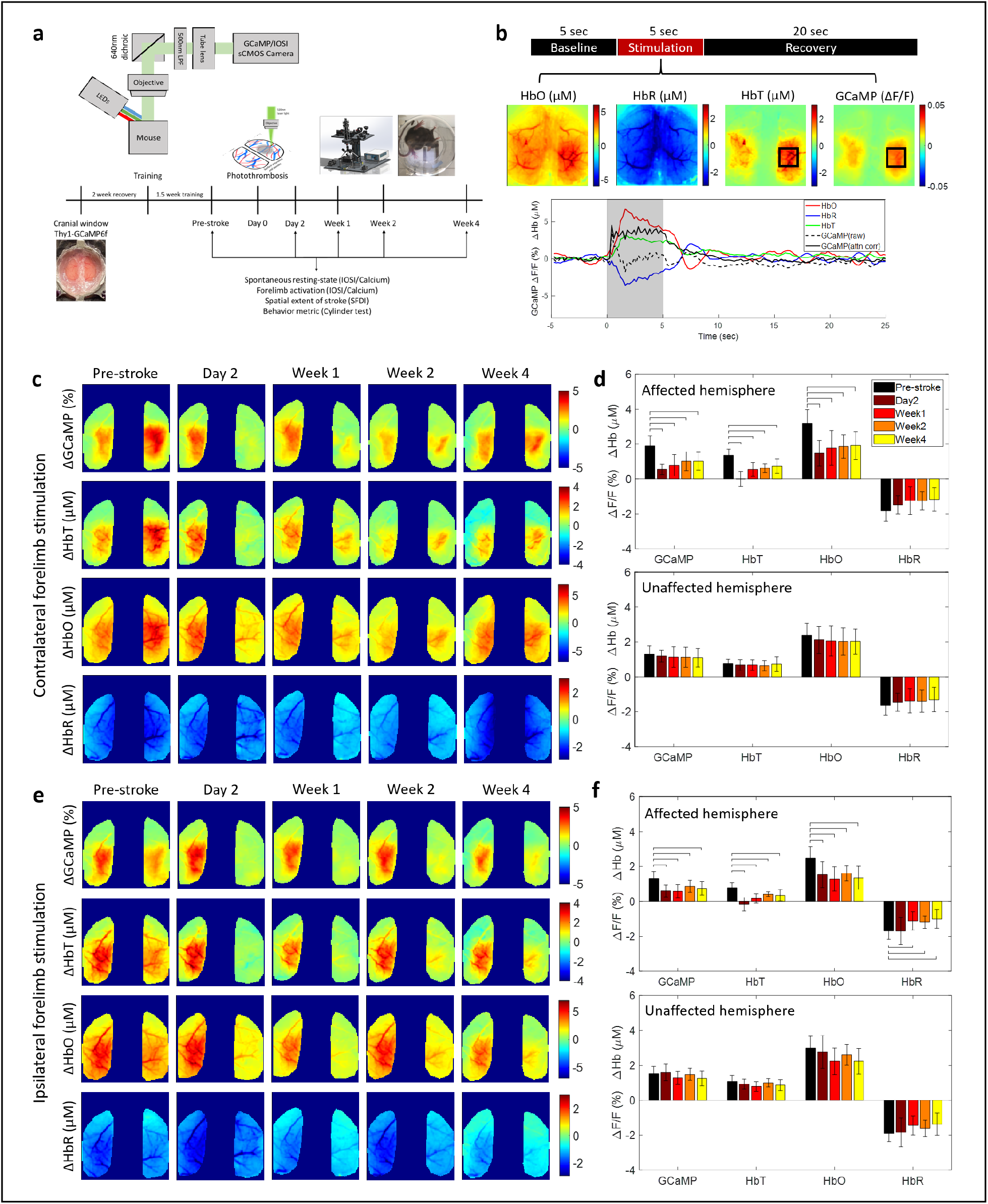
Simultaneous calcium and hemodynamic imaging post-stroke. (a) Simplified imaging schematic and experimental timeline. (b) Top: Block design of each trial in a stimulation session, middle: trial averaged spatial maps of HbO, HbR, HbT, and corrected GCaMP, during 5 sec of air-puff stimulation to the left forelimb, bottom: trial averaged time course of each measurement, note that raw uncorrected GCaMP drops immediately following the rise of the hemodynamic response and corrected GCaMP shows elevated responses through the full stimulation period. (c) Trial-averaged spatial maps of calcium and hemodynamics showing magnitude of the response during 5-sec stimulation of the contralateral (affected) forelimb at each time point before and after stroke in one example mouse. (d) Response magnitudes during affected forelimb stimulation for all mice (n=12) in the affected (top) and unaffected (bottom) hemispheres, histograms are mean ± std. (e) Same as in (c) during stimulation of the ipsilateral (unaffected) forelimb. (f) Same as in (d) during stimulation of the ipsilateral (unaffected) forelimb. Bars in (d) and (f) indicate significance of p<0.05.

### Acute stroke leads to alterations in the correlation between evoked calcium and hemodynamic responses

To evaluate NVC, we examined whether aspects of the observed hemodynamic responses were correlated with the underlying calcium activity during sensory stimulation of the impaired forelimb. Fig 2a shows the trial-averaged mean and standard deviation of the time course of calcium, measured as a percent change in fluorescence (top row), and change in HbO and HbR, measured in µM (bottom row), averaged from all pixels within the affected hemisphere at pre-stroke, 1 week, and 4 weeks post-stroke. Spatial response maps, obtained during the 5-sec stimulation period, were then thresholded at each time point to 75% of the peak response and all pixels that lie above that threshold were used for correlation analysis (Fig 2b). We first examined the similarity in the response areas between the evoked calcium response and hemodynamic measures. Similarity was calculated using the Dice coefficient, which provides a measure of the percent overlap, or union, of two images (Fig 2c). HbT and HbO showed high overlap (60%) with GCaMP before the stroke, indicating that GCaMP and hemodynamic responses were spatially localized, while HbR had a weaker spatial overlap with GCaMP (20%). Within the ipsilesional hemisphere, HbT and HbO showed a significant reduction in the spatial overlap with GCaMP across all time points after stroke, with a larger reduction in the acute time points of day2 and week1 compared to chronic time points. In contract, the overlap between HbR and GCaMP was not significantly altered. Similarity between GCaMP and hemodynamic response maps in the contralesional hemisphere were not significantly altered after stroke (Supplementary Fig 5a).

**Figure 2:**
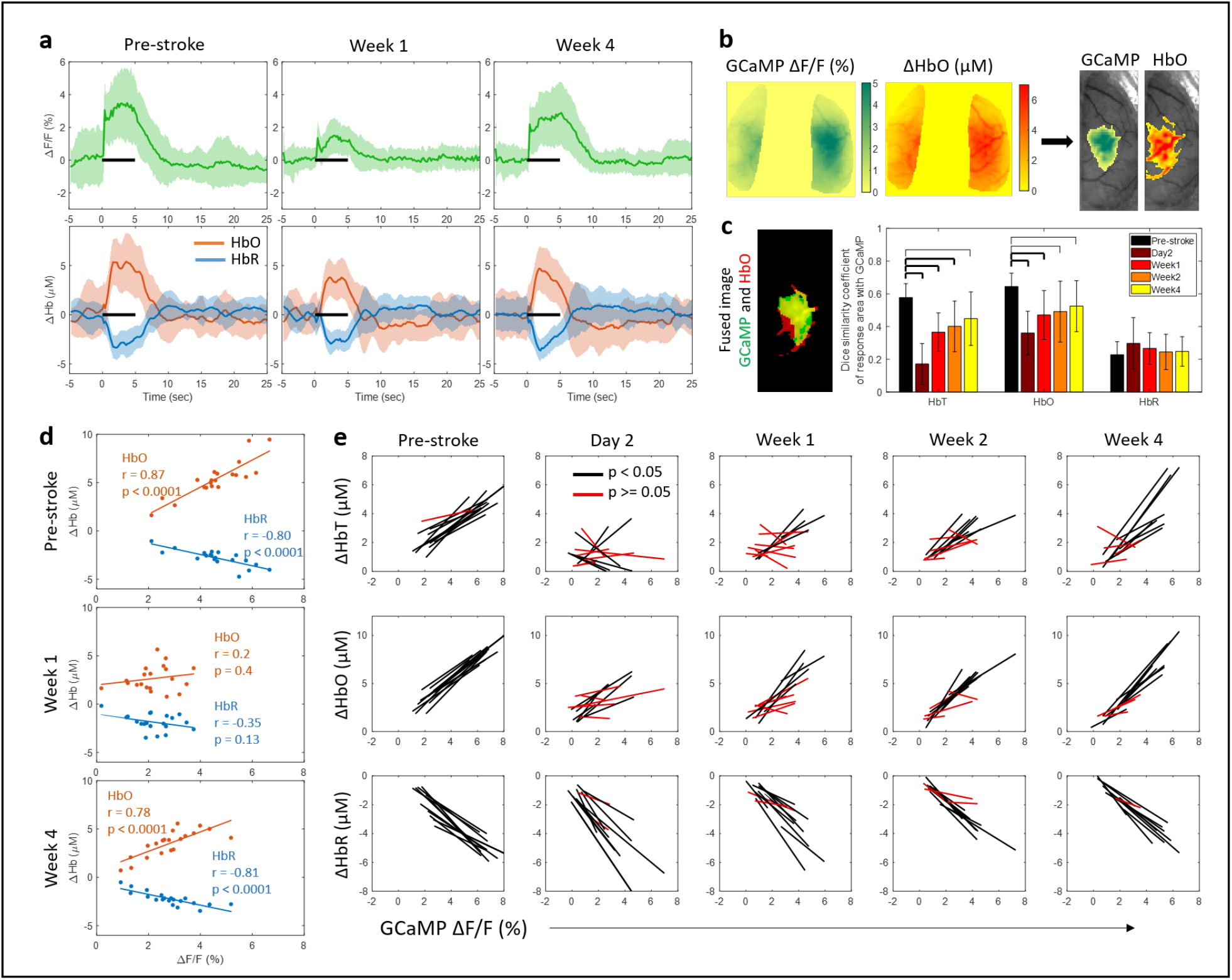
Correlation between evoked calcium and hemodynamic responses. (a) Trial-averaged time-course showing mean (± std) of GCaMP (top) and HbO and HbR (bottom) for all pixels within the affected hemisphere at the pre-stroke baseline, 1 week, and 4 weeks post-stroke. Note the drop in response to stimulation (black bar) at week 1. (b) Threshold algorithm applied to GCaMP and Hb responses. (c) Overlap between the response area of GCaMP and HbO, left: single trial fused image for reference, GCaMP is green, HbO is red, and overlap region is yellow, right: Dice similarity coefficients across all mice (n=12) and time points. Thick bars: p<0.01, thin bars: p<0.05. (d) Correlation of response magnitudes between GCaMP and HbO and HbR for one mouse at pre-stroke, week 1, and week 4. Inset numbers represent correlation value and significance of fit. (e) Correlation of calcium and hemodynamics across all mice (n=12) over all time points; each line represents one mouse. Black lines represent significant correlation and red lines represent no significance.

Next, we calculated the average magnitude of the response during 5 seconds of air-puff stimulation within all pixels above 75% of peak activation. We then correlated the magnitude of the GCaMP response to the magnitude of the HbO and HbR responses for each stimulus trial. Fig 2d shows an example pre- and post-stroke dataset from one mouse; each dot in the scatter plot represents data from one trial within a block of 20 trials. There was high correlation between the evoked GCaMP responses and HbO, as well as GCaMP and HbR, prior to stroke, demonstrating healthy coupling between neural activity and hemodynamics. The correlation was lost 1 week after stroke following a reestablishment by week 4. This evolution of correlation was seen across the cohort of animals (Fig 2e). Both HbT and HbO showed significant loss in correlation with GCaMP in the acute phase, implying that calcium responses in the acute phase were not necessarily represented in the observed hemodynamic response. However, this loss in correlation could also be due to the small amplitudes of the signal, which can result in larger noise and thus low correlation. Additionally, those mice that had a residual loss of correlation at week 4 compared to those that fully recovered, also had a worse correlation between GCaMP and HbT/HbO responses in the acute phase (Supplementary Fig 4). The correlation between calcium and hemodynamic responses in the contralesional hemisphere was preserved throughout the recovery period (Supplementary Fig 5b).

### Acute stroke distorts the shape of the neurovascular response within the peri-infarct zone that is restored in the chronic phase

The next question we asked was whether the shape of stimulus-induced neurovascular response was preserved across the acute and chronic phases of stroke recovery. To that end, we estimated a hemodynamic response function (HRF) (or impulse response function (IRF)), which is the kernel that, when convolved with the GCaMP signal, provides an estimate of the hemodynamic activity. Linear least-squares deconvolution was used to estimate the HRF from the data as established previously^32,37^. First, we validated the method using the baseline (pre-stroke) data. We calculated the HRF using the entire time-course for all pixels that responded to forelimb air-puff stimulation (>75% of peak response) in HbT maps (Fig 3a, top). We observed the expected and characteristic shape of the HRF, a post-stimulus overshoot, peaking at approximately 1 sec following stimulation, followed by an undershoot, as reported previously^32,38^. Next, we applied the same procedure to data collected 2 days following stroke using the same brain region that originally responded to forelimb stimulation. This analysis resulted in a distinctly altered HRF, suggesting a disruption to neurovascular coupling (Fig 3a, bottom). Fig 3b shows the time-course of four individual stimulation trials; the measured GCaMP signal is overlaid with the measured HbT and estimated HbT, where the estimated HbT was obtained by convolving the GCaMP signal with the time-point specific HRF kernel. Pre-stroke, the measured and estimated HbT showed good overlap (Fig 3b, top), while at day 2 after stroke the overlap was poor (Fig 3b, bottom). Additionally, at day 2 after stroke, there was no response to stimulation, and we observed large oscillations in the measured hemodynamic signal. A Pearson’s correlation coefficient was calculated between the measured and estimated HbT signal at each pixel for both hemispheres of the brain (Fig 3c), and we observed high correlation across the somatosensory cortex in both hemispheres before the stroke (Fig 3c, top). From this we can conclude that hemodynamic activity was coupled to the underlying calcium activity prior to stroke. Regions closer to motor cortex showed lower correlation compared to regions within sensory cortex. Higher correlation in the sensory cortex could be due to the presence of air-puff stimulus, which could be driving both calcium and hemodynamic responses and strengthening our observation of neurovascular coupling. This hypothesis could be validated by comparing the HRF and correlation obtained during resting-state and sensory stimulation sessions. Prior work has shown that neural activity is more weakly correlated to hemodynamics during resting state and this can also be validated from our data (Supplementary Fig 6)^39^. Our resting-state data still show a relatively high correlation, which could be due to natural behavior of the mouse, such as whisking and grooming, driving cortical activity. At day 2 after stroke there was a loss in correlation between the measured and predicted HbT as indicated by drop in the Pearson’s correlation coefficient (Fig 3c, bottom).

**Figure 3:**
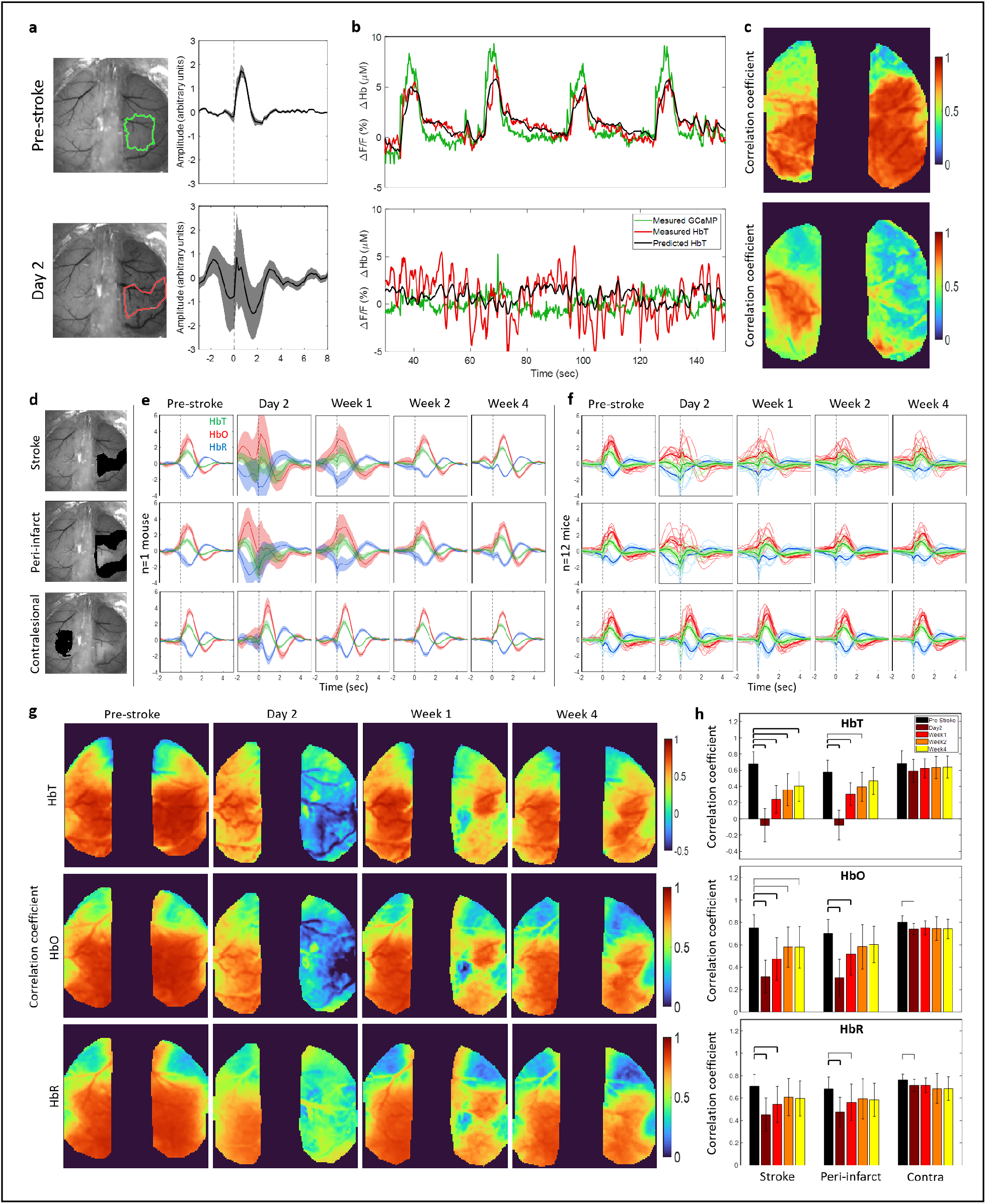
Neurovascular coupling with linear least-squares deconvolution. (a) Hemodynamic response function (HRF) before (top) and 2 days after stroke (bottom) in the forelimb and stroke regions outlined in green and red respectively. (b) Time course of 4 stimulation trials showing measured GCaMP signal overlaid with measured HbT and predicted HbT, obtained by convolving the GCaMP signal with the HRF kernel, at pre-stroke and day 2 for the regions outlined in (a). (c) Pearson’s correlation coefficient for measured HbT and predicted HbT for pre-stroke (top) and 2 days after stroke (bottom). (d) Regions used to extract HRF in (e) and (f). (e) HRF obtained by deconvolution model for HbT, HbO, and HbR, for one example mouse at each time point before and after stroke. Note the deviation in HRF compared to pre-stroke in the acute phase within the stroke and peri-infarct, and a return to pre-stroke HRF at week 4. (f) Same as in (e) for all mice (n=12). Each line represents the HRF for one mouse. (g) Pixel-by-pixel Pearson’s correlation coefficient between measured and predicted HbT (top), HbO (middle), and HbR (bottom). Predicted HbX is obtained by convolving the GCaMP signal at each time point with a mean HRF obtained from pre-stroke data. (h) Pearson’s correlation coefficient quantified across all mice within the stroke core, peri-infarct, and contralesional forelimb region. Thick bars: p<0.01, thin bars: p<0.05. Note the sustained reduction of correlation coefficient within the stroke core but recovery within the peri-infarct for HbT and HbO.

The same convolution model was applied to calcium and hemodynamic data to estimate the HRF post-stroke across all animals and time points. The post-stroke HRF was calculated for the stroke core, the peri-infarct region, which included all the pixels within 0.5 mm from the stroke core boundary, and the contralesional forelimb region. The HRF was also calculated for HbO and HbR in addition to HbT. Fig 3d shows an example mouse where the stroke core, peri-infarct, and contralesional forelimb are highlighted in black. We then followed the evolution of the HRF for each hemodynamic measure after stroke. Fig 3e shows the mean and standard deviation of the HRF for one example mouse. We observed a significant deviation in the HRF within the stroke core and peri-infarct region in the acute phase of recovery. Following the acute phase, the chronic phase showed a recovery in the HRF. The contralesional HRF remained largely unaffected by the stroke. Similar trends were observed across all animals (Fig 3f), where acute stroke resulted in deviation of the HRF in the core and peri-infarct, while the contralesional hemisphere was unaffected. In the chronic phase the HRF showed better recovery of the shape, with respect to pre-stroke HRF, in the peri-infarct region. The HRF within the stroke core continued to show deviation in some animals.

Here, we describe deviation of the HRF in terms of the qualitative similarity of shape to the pre-stroke HRF. However, even if the shape of the HRF is different, it could still be used to accurately predict hemodynamics. Therefore, we next tested the ability of the HRF at each time point to predict hemodynamics. Supplementary Fig 7a shows a pixel-by-pixel map of the Pearson’s correlation coefficient for one example mouse for all three hemodynamic measures before and after stroke. We observed a clear drop in correlation coefficient within the affected hemisphere, specifically in day 2. Over time, through the recovery period, we observed some return of correlation between the measured and predicted hemodynamics. The correlation coefficient was quantified across all mice in the stroke core, peri-infarct, and contralesional forelimb region (Supplementary Fig 7b). The stroke core showed a significant reduction in correlation coefficient across all time points compared to before stroke, implying that the hemodynamic response captured the underlying neural activity significantly worse compared to pre-stroke. Additionally, this shows that the deviation in shape of the HRF was also associated with a lack of correlation between measured and predicted hemodynamics. There was also a significant decrease in the ability of the neurovascular coupling model to capture the hemodynamics from the measured GCaMP signal within the peri-infarct region in the acute phase of day 2 and week 1. However, unlike the stroke core, the peri-infarct showed recovery in terms of reestablishing the correlation between the measured and predicted hemodynamics in the chronic phase, which was also associated with a return of the HRF shape to the pre-stroke shape.

From the shape of the HRF we can clearly see that the neurovascular coupling model is not behaving as expected during day 2 and week 1. Most notably, we see that the HRF is not flat prior to stimulus onset at time = 0 as we would expect. We have provided more flexibility in our model by allowing it to use GCaMP events that have not happened yet to find the best fit. In the pre-stroke case this negative time region is a flat line at zero indicating that future GCaMP events have no influence on the current hemodynamics, as expected. However, after stroke, specifically at day 2 and week 1, the HRF is no longer flat before time zero. While it is physiologically not possible for future GCaMP events to influence current hemodynamics, this deviation in the HRF indicates that there are possibly additional dynamics that are not captured by the original model and the model is just trying to find the best fit with the given data. We can overcome this limitation and test deviations in neurovascular coupling by testing how well we are able to predict the post-stroke hemodynamics with the pre-stroke HRF, since we know that the pre-stroke HRF is behaving as expected. To test this, we calculated the mean HRF for each mouse from pre-stroke “healthy” data and convolved it with the post-stroke GCaMP time-course and obtained the correlation with this predicted and measured hemodynamics (Fig 3g). Similar to when we used the time-point specific HRF, there was a significant drop in correlation within the stroke and per-infarct regions in the acute phase and a recovery within the peri-infarct region in the chronic phase when using the pre-stroke “healthy” HRF (Fig 3h). Unlike the time-point specific HRF correlations (Supplementary Fig 7a, 7b), the healthy HRF correlations with post-stroke data showed virtually no correlation between the measured and predicted HbT and only a small correlation in HbO at day 2. This suggests that the neurovascular coupling model described for healthy brains is not sufficient to describe post-stroke neurovascular dynamics during the acute phase. The stroke core continued to show poor correlation even in the chronic phase at week 4 but the peri-infarct region exhibited a recovery.

### Acute stroke leads to increases in power of global brain oscillations

Stroke is known to have a profound effect not only on the local network but also on the contralesional and subcortical networks of the brain. Additionally, in our neurovascular coupling analysis we observed an increase in oscillatory dynamics in the hemodynamic signal. Through our wide-field imaging approach we can assess the effect of stroke on both hemispheres of the brain during resting-state. To assess brain-wide variations in the signals we first investigated the overall change to signal patterns. Fig 4a shows the resting-state time-courses of GCaMP and HbO signals at pre-stroke and 2 days post-stroke within the ipsilesional peri-infarct (Fig 4a, top) and the contralesional forelimb regions (Fig 4a, bottom) that was filtered at 0.009-0.4Hz, which covers the low and high frequency hemodynamic signal ranges used in prior work^40^. A feature of note here is the increase in amplitude of the HbO signal at day 2 in the ipsilesional hemisphere (light red line in Fig 4a top) compared to pre-stroke, but an increase in amplitude of both the HbO and GCaMP signal at day 2 within the contralesional hemisphere. We validated this increase in amplitude by calculating the variance in the overall signal (Fig 4b). GCaMP showed only minor alterations in variance while HbO showed a large increase in the variance of its signal at day 2, which was resolved by week 4. To address whether this increase in the amplitude of the signal was an increase in the power of the signal across all frequencies or specific to a particular frequency. We calculated the power spectrum of the GCaMP and hemodynamic signal within the affected and unaffected hemisphere (Fig 4c). There was an overall increase in power across all frequencies at 2 days after stroke in the HbO signal of the ipsilesional hemisphere. Moreover, there was a significant increase in power of the hemodynamic signal at 2 days and 1 week after stroke at specifically around 0.25 Hz within the ipsilesional hemisphere. The contralesional hemisphere on the other hand showed increased power at 0.25 Hz at day 2 after stroke in both GCaMP and hemodynamics. Fig 4d shows the area under the curve in the frequency range of 0.1-0.3 Hz, where the largest increase in power was observed. This increase in power at 0.25 Hz, which is typically higher than normal for hemodynamics, could be a result of increased vasomotion. Evidence from prior work in human laser doppler flowmetry and magnetoencephalography (MEG) has suggested that stroke affected arterioles showed elevated power^41,42^.

**Figure 4:**
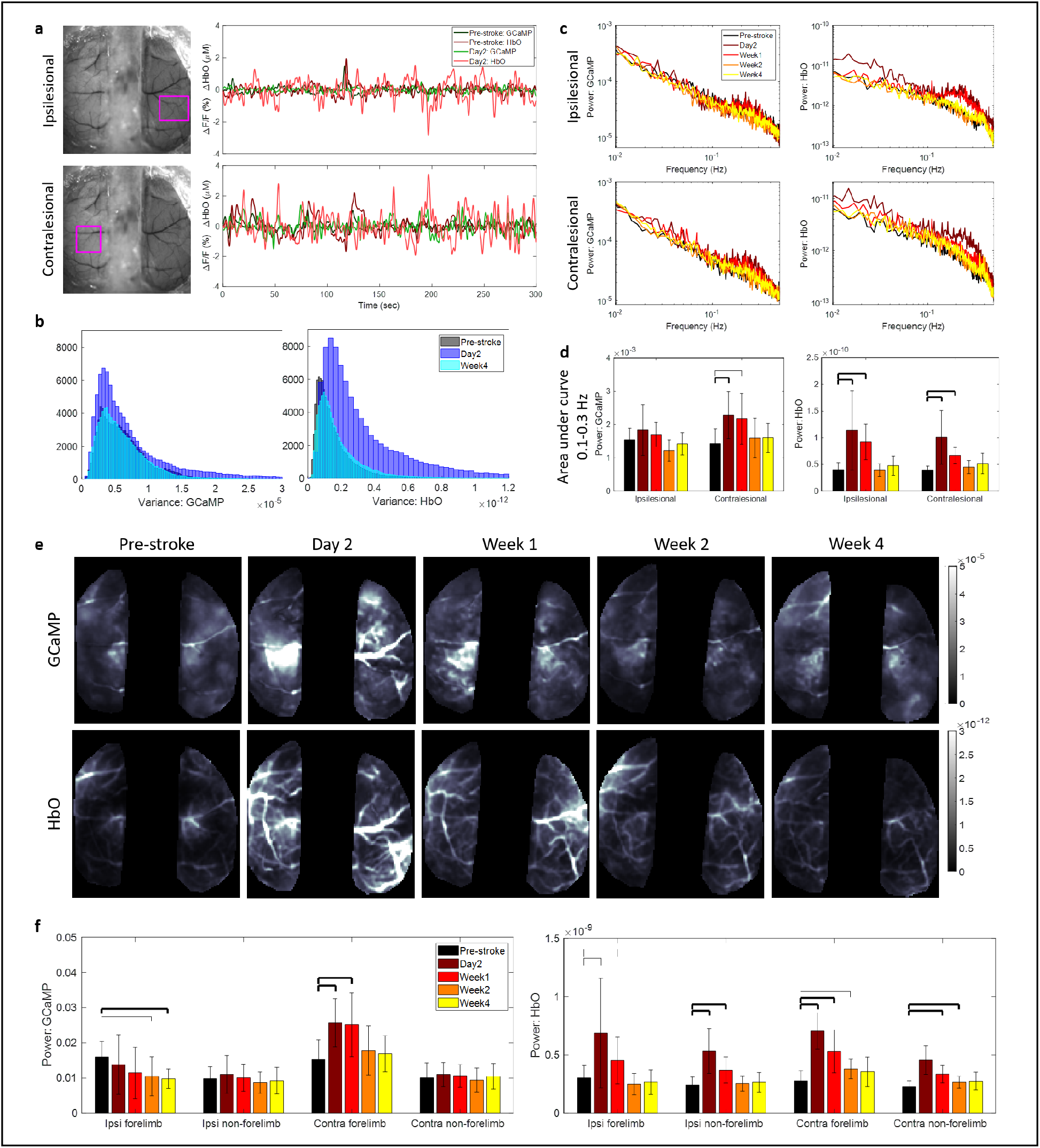
Global brain oscillations following stroke. (a) Raw time traces of filtered (0.009-0.4 Hz) calcium and hemodynamic signals before and 2 days after stroke within the ipsilesional (top) and contralesional (bottom) hemispheres in ROI marked with pink box. Note the increase in amplitude of HbO in both hemispheres at day 2 and increase in GCaMP amplitude only in the contralesional hemisphere. (b) Histogram of variance in the mean signal, after global signal regression, for GCaMP (left) and HbO (right) at pre-stroke, day 2, and week 4. (c) Frequency spectrum of the power of the GCaMP (left) and HbO (right) signal in the ipsilesional (top) and contralesional (bottom) hemispheres. (d) Area under the curve within 0.1-0.3 Hz frequency band. Thick bars: p<0.01, thin bars: p<0.05. (e) Spatial maps of average power across 0.009-0.4 Hz frequency band for GCaMP (top) and HbO (bottom) at each time point. (f) Mean power assessed in each hemisphere within the forelimb and non-forelimb areas. Thick bars: p<0.01, thin bars: p<0.05.

We then asked if this increase in power of the GCaMP, in the contralesional hemisphere, and hemodynamic signal, in both hemispheres, was uniform across the hemispheres or specific to any distinct brain region. Fig 4e shows spatial maps of the average power for GCaMP and HbO for one typical mouse. We clearly see increased power in GCaMP in the contralesional hemisphere and increased overall power in HbO at day2 and week 1 compared to pre-stroke. Surprisingly, the increase in GCaMP power appeared specific to the contralesional forelimb region, while the power increase in HbO appeared global. This was validated across all mice (Fig 4f), which showed that there was a significant increase in power within only the contralesional forelimb and not the rest of the contralesional hemisphere. The HbO signal on the other hand showed increases across all regions, the ipsilesional and contralesional forelimb and non-forelimb areas. There was also a decrease in the GCaMP signal within the ipsilesional forelimb region in the chronic phase, which is likely due to loss of neurons within that region.

### Photothrombotic stroke disrupts resting state interhemispheric functional connectivity only in the very acute phase

Stroke has also been known to affect functional connectivity across large scale brain networks^43,44^. To address the possibly differential effects of stroke on calcium and hemodynamic global brain dynamics we asked whether resting state functional connectivity (RSFC) showed similar dynamics during the recovery phase. Prior work in healthy animals has showed that at low (0.009-0.08 Hz) and high (0.08-0.4 Hz) frequency bands, which are typically used in BOLD fMRI and functional connectivity IOSI studies, functional connectivity structures between GCaMP and HbO were in high agreement^33^. But as applications of hemodynamic RSFC are extended into the stroke field it is not only important to understand the underlying physiology that those signals represent but also what aspects of connectivity are altered and are sensitive measures for the stroke^29,45^.

To that end, we looked at various aspects of RSFC in the low and high frequency bands across the GCaMP and HbO maps. First, we assessed connectivity of the ipsilesional forelimb area to the contralesional hemisphere (Fig 5a). In healthy pre-stroke animals, seed-based forelimb connectivity maps were consistently normal when compared to prior work, while acute stroke showed alterations in forelimb connectivity to the contralesional hemisphere^45–47^. Fig 5a(i) shows the forelimb connectivity maps for GCaMP at pre-stroke, day2, and week4 in the low frequency band. We then quantified the differences between pre-stroke and each post-stroke time point by calculating the proportional area of the cortex above a certain correlation coefficient threshold that ranged from 0 to 0.9 (Fig 5a(ii)). A slight decrease in connectivity was observed in the GCaMP map and a large decrease was observed in HbO at 2 days post-stroke (Fig 5a(ii), Supplementary Fig 8a). HbO continued to show reduced forelimb connectivity at all time points after stroke at specific thresholds (Supplementary Fig 8a), however, GCaMP connectivity appeared largely restored at later time points. A Dice similarity index was calculated between the GCaMP and HbO maps across all thresholds, which showed a deviation in similarity only at day2 after stroke, while maps were consistent at all other time points (Fig 5a(iii), Supplementary Fig 5a). A similar approach was used for calculating interhemispheric connectivity, global connectivity, and contralesional forelimb intrahemispheric connectivity as well as all measures in the higher frequency band. Trends across time points and thresholds for all measures were largely similar in both frequency bands. There was a significant drop in interhemispheric connectivity at 2 days in both GCaMP and HbO, which was restored at later time points in GCaMP but continued to persist, to a lesser extent, in HbO until week1 (Fig 5b,f). Surprisingly there was a small but significant increase in global connectivity at day2 in GCaMP (Fig 5c,g, Supplementary Fig 8c). Spontaneous recovery over four weeks resulted in reestablishment of global connectivity networks in both GCaMP and HbO. Since we observed increase in the calcium power within the contralesional hemisphere (previous section) we also asked whether contralesional forelimb connectivity was altered. We observed a significant increase in contralesional forelimb connectivity within the contralesional hemisphere at day 2 after stroke (Fig 5d,e). This suggests that increases in the power of the calcium signal within the contralesional forelimb was associated with an increase in its functional connectivity to other regions of the brain. The increase observed in the global connectivity index could be due to this increased connectivity of the contralesional forelimb.

**Figure 5:**
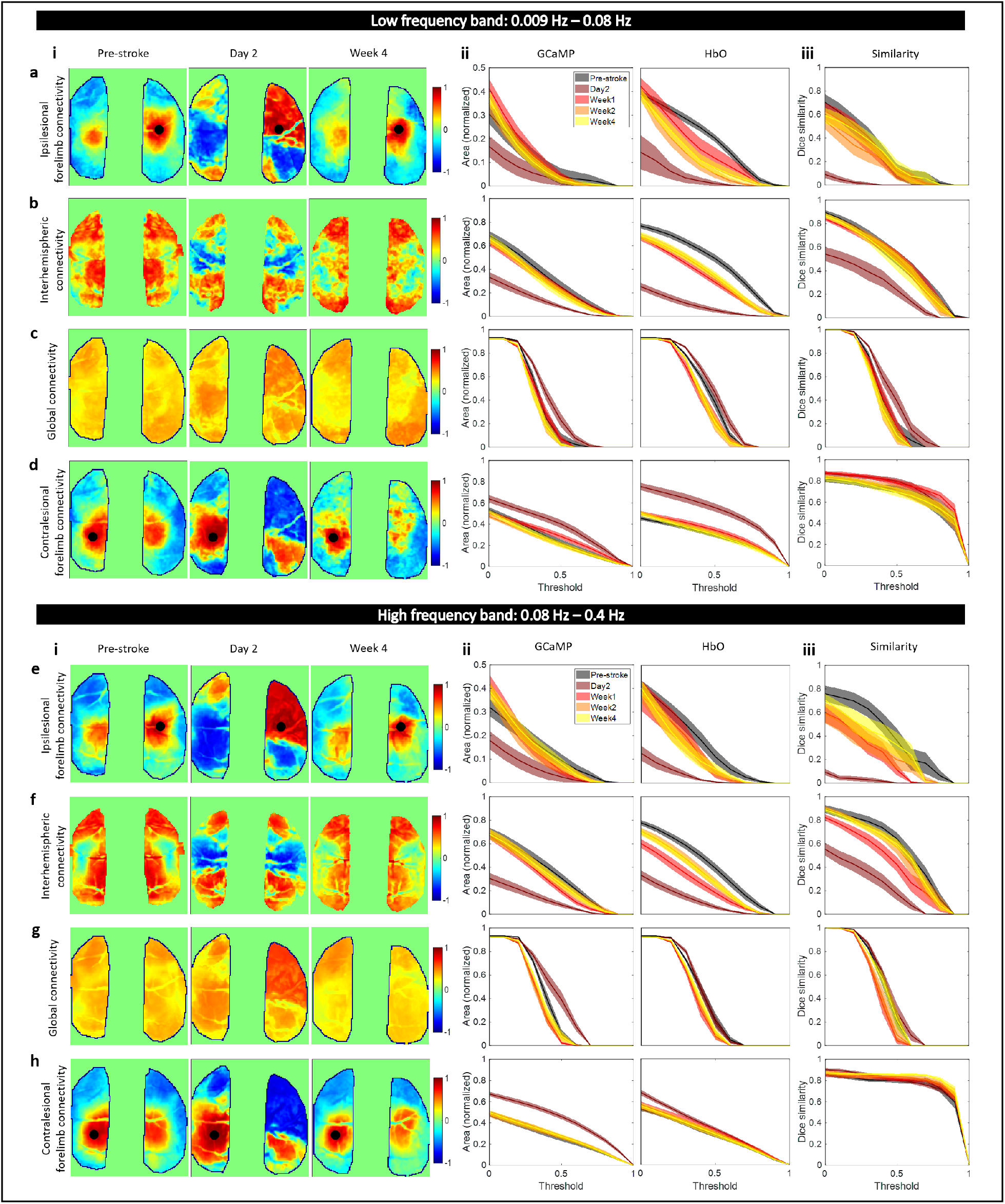
Global brain network dynamics assessed with RSFC. Spatial maps of ipsilesional forelimb connectivity (a(i),e(i)), interhemispheric connectivity (b(i),f(i)), global connectivity (c(i),g(i)), and contralesional forelimb connectivity (d,h) in the low frequency band (a,b,c,d) and the high frequency band (e,f,g,h) at pre-stroke, day 2, and week 4. Proportional area of cortex over threshold for GCaMP and HbO at each time point for ipsilesional forelimb connectivity (a(ii),e(ii)), interhemispheric connectivity (b(ii),f(ii)), global connectivity (c(ii),g(ii)), and contralesional forelimb connectivity (d(ii),h(ii)) in the low frequency band (a,b,c,d) and in the high frequency band (e,f,g,h). Dice similarity coefficient for overlap between area covered by GCaMP and HbO for ipsilesional forelimb connectivity (a(iii),e(iii)), interhemispheric connectivity (b(iii),f(iii)), global connectivity (c(iii),g(iii)), and contralesional forelimb connectivity (d(iii),h(iii)) at all time points in the low frequency band (a,b,c,d) and the high frequency band (a,b,c,d).

From these data we extrapolate that connectivity of both the impaired and unimpaired forelimb and interhemispheric connectivity for both GCaMP and HbO were reliable measures to indicate stroke, given our photothrombotic model, at day 2. Disruptions to interhemispheric connectivity persisted until week 1 after stroke, however other metrics assessed were indistinguishable from pre-stroke. While global connectivity provides a concise method as a seed-independent approach of functional connectivity, in our case it was a weaker metric for following the stroke recovery process.

### Correlating acute phase cortical metrics to long-term behavior outcomes

To enable translation of the cortical measures investigated in this work to potentially clinically relevant outcomes, we measured forelimb performance through the cylinder asymmetry test before stroke and at each imaging time point after stroke. Photothrombotic stroke to the forelimb somatosensory area led to deficits in the use of the impaired forelimb (Fig 6a). Mice used their impaired forelimb 50% less than baseline in the first week following stroke, however, over time with spontaneous recovery, mice showed a significant increase in the use of the impaired forelimb by week 4 compared to day 2 (Fig 6a). An important factor in human strokes that is often missed in animal models is the variability in the extent of damage and impairment caused by the stroke. The extent and location of the damage due to stroke as well as the early spontaneous recovery mechanisms play a significant role long-term outcome^3,4,48^. A number of early biomarkers that might have potential as indicators of behavioral outcome are under active investigation in both humans and in animal studies^49,50^. While it would be ideal to introduce a controlled level of variability into animal models to study variable recovery and to identify biomarkers that indicate recovery, such a method of stroke induction does not yet exist that also meets all the other criteria for a physiological stroke, such as preventing the use of anesthesia during stroke induction. Our optimized photothrombotic model introduces uncontrolled variability that mimics human variability to some extent and allows correlation of behavioral outcome to cortical biomarkers. The right panel of Fig 6a shows the extent of recovery in forelimb asymmetry for individual mice at week 4. In this section we outline how cortical measures obtained in all the previous sections correlate to these variable long-term behavioral outcomes.

**Figure 6:**
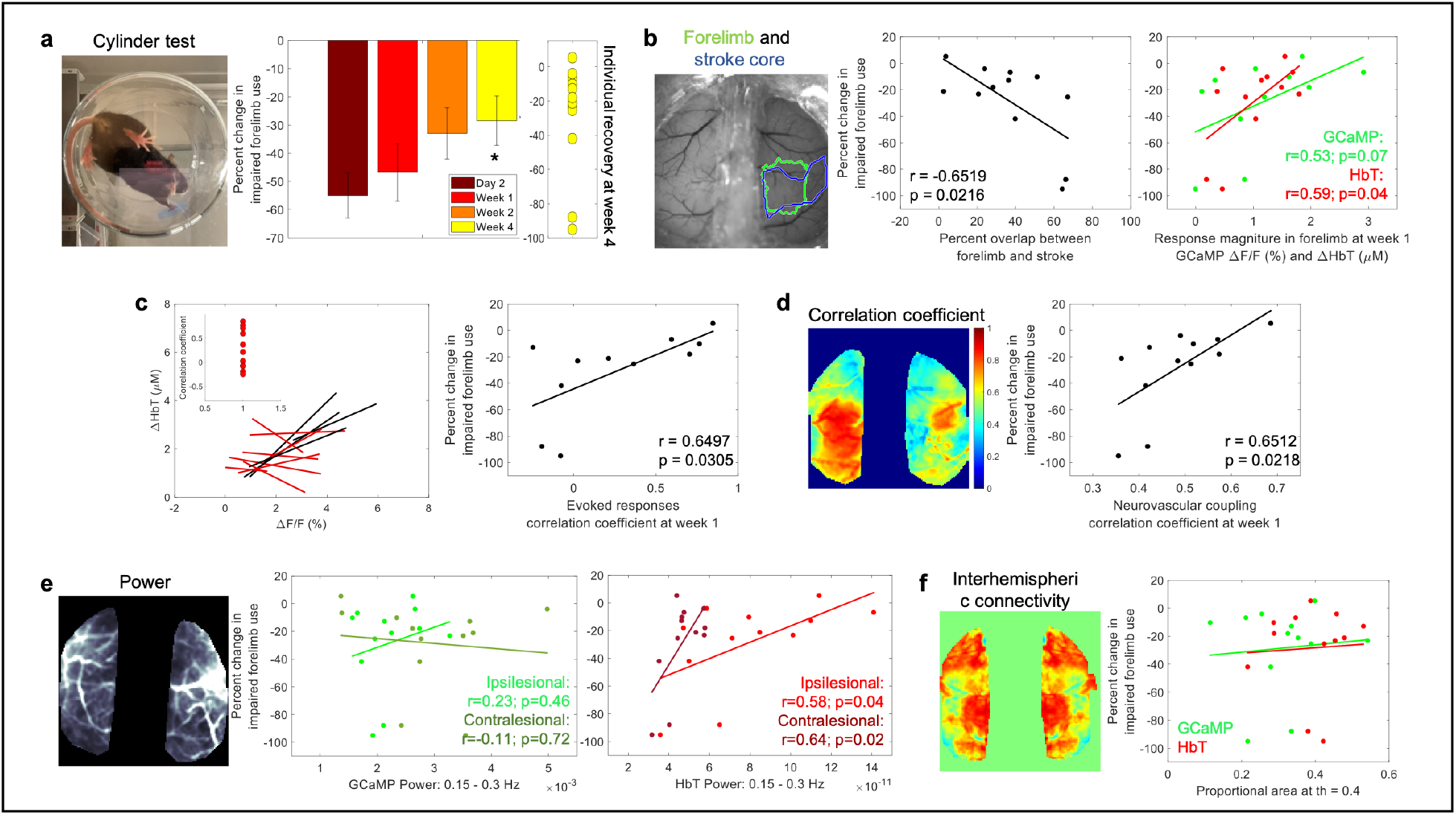
Correlating cortical metrics to behavior outcomes. (a) Forelimb asymmetry, assessed with the cylinder test, calculated as a change in impaired forelimb use from pre-stroke, right: recovery of individual mice at week 4. (b) Left: reference image showing outlines of pre-stroke forelimb region and stroke core at 1 week, middle: correlation between overlap of forelimb and stroke with forelimb asymmetry at week 4, right: correlation between response magnitude at week 1 for GCaMP and HbT with forelimb asymmetry at week 4. (c) Left: correlation of evoked responses of GCaMP and HbT, right: correlation between the correlation coefficient of evoked responses at week 1 and forelimb asymmetry at week 4. (d) Left: correlation coefficient between measured HbT and HbT predicted by convolving GCaMP and IRF, right: correlation between neurovascular coupling correlation coefficient at week 1 and forelimb asymmetry at week 4. (e) Correlation between power of GCaMP and HbT in frequency band 0.15-0.3 Hz in the ipsilesional and contralesional hemispheres and forelimb asymmetry at week 4. (f) Correlation between resting state interhemispheric connectivity and forelimb asymmetry at week 4.

First, since the cylinder test is sensitive to forelimb use, we tested whether the extent of forelimb area that was damaged due to stroke predicted behavioral outcomes. We calculated the percent overlap between the pre-stroke forelimb region and the stroke outline obtained at 1 week after stroke from SFDI (Fig 6b). There was a significant negative correlation indicating that worse behavior outcomes correlated with a larger portion of the forelimb being damaged by the stroke. We next asked whether functionality in the surviving portion of the forelimb region promoted behavioral recovery. Here, we calculated the magnitude of responses within the original forelimb region for GCaMP and hemodynamics at week 1. Both GCaMP and hemodynamics showed trends towards a positive correlation between response magnitude and better outcomes, with only the HbT showing a significant correlation (Fig 6b). Next, we assessed the relationship between acute neurovascular coupling and behavior outcomes. This was calculated in two ways, using the correlation between GCaMP and HbT responses and using HRF. The correlation coefficients obtained from the magnitude of evoked responses between GCaMP and HbT at week 1 during forelimb stimulation significantly correlated with behavioral outcomes (Fig 6c, data from Fig 2). The correlation coefficients obtained from the neurovascular coupling HRF model between the measured and estimated HbT at week 1 also showed a significant correlation with behavioral outcomes implying that preserved or improved neurovascular coupling at week 1 might be indicative of better long-term recovery (Fig 6d).

We performed similar calculations with the global brain metrics of power of the signal and interhemispheric connectivity from RSFC. Average power was calculated within the narrow frequency band (0.15–0.3Hz) for both GCaMP and hemodynamics and separated into ipsilesional and contralesional hemispheres. While GCaMP did not show any trends with behavior, acute hemodynamic oscillations showed strong positive trends with behavior outcomes (Fig 6e). Specifically, increased power in the HbT signal of the contralesional and ipsilesional hemispheres in the acute phase of stroke significantly correlated with behavior outcomes. Interhemispheric connectivity at week 1, or any other RSFC metric, did not show any correlations with long-term behavior outcomes, further implying that RSFC might not be a sensitive metric for targeted photothrombosis. Overall, we have identified several cortical metrics within the acute phase of stroke recovery that had the potential to delineate animals that tend to show better spontaneous recovery versus animals that had poorer recovery.

## Discussion

While functional neuroimaging has great potential for treating and monitoring patients in the acute and chronic phases of stroke recovery, the interpretations of these signals and their reliability as a neural correlate is still under active investigation. In this study, we used an animal model of stroke, which was optimized for high clinical relevance, to investigate the relationships between neural activity, assessed with a fluorescent calcium indicator, and cerebral blood volume, assessed with changes in oxy and deoxy hemoglobin, during longitudinal stroke recovery. We showed that acute stroke leads to disruptions in neurovascular coupling, which is restored in the chronic phase. Neurovascular uncoupling was primarily experienced within the affected hemisphere and early recoupling and recovery of cortical function within the preserved forelimb region and peri-infarct zone was an indicator of better recovery. Additionally, we showed that acute stroke leads to increases in global brain oscillations, which show distinct spatial characteristics in GCaMP and hemodynamics.

The results from this study have several implications for the interpretations of hemodynamic signals in terms of the underlying physiology in both pre- and post-stroke. In the healthy brain we showed that with simultaneous multi-modal imaging of neural calcium activity and hemodynamics we can track subtle differences in sensory evoked response dynamics on a trial-by-trial basis. This allowed us to correlate intra-animal changes to evoked responses across the cohort and at each time point after stroke. We correlated each animal’s responses individually due to the variability in the extent of ischemic damage among animals introduced by our stroke model. This allowed us to track the changes in each animal individually and we found that there was a significant loss in correlation between evoked calcium and hemodynamic responses in the acute phase at day 2 and week 1. Correlation was reestablished in most animals by week 4 signifying spontaneous recovery and improved behavior. A small number of animals continued to show loss of correlation between evoked calcium and hemodynamic responses across both the acute and chronic time points and these animals were associated with poor behavior outcomes. There was also a significant positive trend between correlation of evoked responses, specifically between calcium and HbT, at week 1 and behavior outcomes at week 4 across all mice. Taken together with the significant correlation between early HbT response magnitudes within preserved forelimb and long-term behavior, this implies that early recovery of hemodynamic responses, HbT in particular, might be indicative of better outcomes.

While correlations of evoked calcium and hemodynamic response magnitudes allowed us to draw conclusions about the similarity, or dissimilarity in the case of stroke, between the two measures, it does not contain quantitative information about their relationship. To quantitatively describe neurovascular coupling, we predicted hemodynamics from calcium activity using linear least-squares deconvolution as had been done previously^32,38^. Similar to previous reports, the measured calcium signal convolved with a calculated HRF kernel predicted the hemodynamic signal to a high degree in healthy animals. There was a higher correlation within sensory regions of both hemispheres compared to more frontal or posterior regions, likely due to sensory stimulation driving cortical activity within somatosensory cortex and strengthening the observed neurovascular coupling signal. The characteristic shape of the HRF was altered after acute stroke, which also corresponded with a significant decrease in the ability of the model to predict hemodynamics within the affected hemisphere. The correlation when using the pre-stroke “healthy” HRF was significantly lower than the correlation when using the time-point specific post-stroke HRF. This indicates that while the model post-stroke was finding the best fit, the resulting HRF was not necessarily similar to the expected neurovascular coupling model under healthy conditions. Using the expected neurovascular coupling model yielded significantly worse correlations. These results suggest that the neurovascular coupling model established in healthy animals was not representative of post-stroke acute phase dynamics and that the observed hemodynamic response is not an accurate representation of the underlying physiology since the HRF was unable to predict hemodynamics accurately during the acute phase. However, it must also be noted that the model assumption of a linear relationship might not hold true after stroke, and the hemodynamic response might be better predicted with an altered non-linear model. Nevertheless, we see restoration of neurovascular coupling, in accordance with the linear model, in the chronic phase of recovery. We observed reestablishment of the expected HRF shape and improvement in the ability of the model to predict hemodynamics, specifically in the peri-infarct region. This suggests that functional neuroimaging might be faithfully representing the underlying neurophysiology in the chronic phase.

In addition to local changes to evoked responses and neurovascular coupling alterations within the affected hemisphere, stroke is known to have a profound impact on global cortical network dynamics such as contralateral and subcortical connectivity^44^. We found that there was an increase in the overall power of cortical signals in both calcium and HbT in the acute phase, which was resolved in the chronic phase. The increase in power of the calcium signal appeared to be specific to the contralesional forelimb region, while the increase in hemodynamic power was global across all vessels and both hemispheres. A prior study conducted with laser doppler flowmetry showed increased oscillations within stroke affected arterioles and suggested increased vasomotion as the cause^42^. Other studies have also showed increases in brain oscillations in stroke and traumatic brain injury^41,51,52^. Vasomotion, which is the oscillating tone of blood vessels independent of heart rate or breathing, is tightly regulated, and maintained by various compartments of the neurovascular unit^53,54^. Vascular autoregulation is impaired after stroke and ionic imbalances in neural, astroglial, and endothelial cells could result in dysregulation of vasoactive molecules and ions and therefore vascular tone^17,18^. On the other hand, we also observed increases in power of GCaMP in the contralesional forelimb. Prior work has shown that stroke leads to increases in brain excitability and disruption of the interhemispheric inhibition through the corpus callosum^55–57^. This could reduce the inhibitory effects that the two hemispheres exert on each other, which could increase excitability within the contralesional hemisphere. There is also evidence of thalamic disinhibition within minutes of ischemic stroke that can unmask ipsilateral pathways^58^. The excitability of thalamocortical pathways contralateral to the stroke may be enhanced because of downregulation on interhemispheric thalamic inhibition. Surprisingly, we found that increased power in the hemodynamic signal in the contralesional hemisphere during the acute phase was correlated with improved behavior outcomes. Prior work has shown that stimulation of activity within the gamma frequency band improved cerebral blood flow, decreased infarct volume, and improved motor behavior, suggesting that modulation of cortical oscillatory dynamics may serve as a target for neuroprotection^59^. Other studies have also shown that increased brain oscillations and excitability promoted recovery in stroke as well as other neurological disorders and suggest its possible use as a biomarker for recovery^51,55,56,60^. A meta-analysis on activation data from over 50 neuroimaging experiments have shown enhanced activity in the homotopic region of the contralesional hemisphere in the acute phase after stroke^61,62^. This enhanced activity appears as spontaneous and synchronous neural activity and has been shown to be a signal for axonal sprouting and reorganization^63^. Taken together with this evidence, we could hypothesize that spontaneous increases in power that we observed in hemodynamic activity might play a role in promoting recovery mechanisms. These oscillations are possibly driven by underlying neural activity at frequencies higher than we can measure with GCaMP, which we are unable to capture due to the slow calcium dynamics compared to neural firing.

A growing number of studies are now using RSFC to assess spatiotemporal correlations in spontaneous hemodynamic signals across different brain regions in healthy and diseased states. In the healthy brain, hemodynamic signals have been found to be bilaterally correlated and synchronized temporally in functionally distinct brain regions and represent the connectivity of underlying intrinsic neural fluctuations^32,46,64^. RSFC has also been used as a sensitive assay to monitor progression of stroke and hemorrhage with the assumption that the altered connectivity represents the altered neural state^29,45^. In this study we show that RSFC of spontaneous calcium activity and hemodynamics show similar trends after stroke, validating prior assumptions. Forelimb and interhemispheric connectivity were disrupted significantly in the very early acute phase and was resolved within week 1 in both calcium and hemodynamics. Moreover, we found that RSFC measures were not predictive of behavioral outcome. This could be because global brain connectivity is more robust to small strokes caused by targeted photothrombosis to the forelimb. A prior RSFC study also showed that somatosensory connectivity was not predictive of behavior but motor and retrosplenial cortices might be better predictors^29^. Due to our window preparation procedure and headbar design for multimodal optical access we were limited in the field-of-view to mainly the somatosensory region and were unable to capture connectivity to other brain regions to their full extent. It is also possible that more sensitive analyses are needed for RSFC to serve as a metric for stroke outcome. We also tested whether increases in power of the calcium activity within the contralesional forepaw was associated with increased functional connectivity through RSFC. We found that intra-contralesional hemisphere connectivity was significantly increased at day 2. This suggests that increased excitability within the contralesional forepaw might result in its increased functional connectivity to surrounding regions as well as the ipsilesional peri-infarct, as seen from the spatial maps of connectivity. Further investigation is needed to understand the link between excitability and functional connectivity and its impact on recovery.

An important factor to note in our study is that we measure calcium dynamics from only excitatory cells. We know, from decades of prior work, that both excitatory and inhibitory cells have important and distinct roles to play in maintaining cortical balance^65^. Additionally, a number of other cell types, such as astrocytes and pericytes, and modulators are involved in regulating blood flow to meet the metabolic demands of the brain^14,66,67^. We also know that these different cell types are impacted differently after stroke^1,2,68^. While the current study used mice with labelled excitatory neurons, the same imaging platform and experimental design can be used to investigate the contributions of other cell types, such as inhibitory cells and glia, to alterations in neurovascular coupling after stroke. Additionally, calcium dynamics assessed with GCaMP6f has been validated to be a reliable measure of neural activity, however, it is still not a direct measure of neural electrical activity. Fast neural dynamics or sub-threshold dynamics may be missed in calcium imaging since the dynamics of calcium are much slower than action potentials or local field potentials. Although performing similar experiments while capturing local field potentials would allow us to assess neural activity directly and provide a higher temporal resolution, we do not believe that using GCaMP has affected our assessment of neurovascular coupling as all our experiments are performed at a temporal resolution higher than what is needed for hemodynamics assessment.

In summary, by simultaneously capturing changes in neural calcium activity and hemodynamics we have assessed various aspects of neurovascular coupling during the acute and chronic phases of stroke recovery. Our data suggest that acute stroke leads to neurovascular uncoupling, implying that functional neuroimaging by fMRI and fNIRS might not accurately represent the underlying neural activity and one needs to use caution when interpreting the results. Neurovascular coupling is restored in the chronic phase, suggesting that these functional neuroimaging methods more faithfully represent the underlying neural activity chronically. Moreover, early recovery of neurovascular coupling and increased power of brain oscillations were predictors of better long-term behavioral outcomes.

## Methods

### Experimental design

All experiments and animal procedures were approved by the Boston University Institutional Animal Care and Use Committee and were conducted following the Guide for the Care and Use of Laboratory Animals. All animals used in this study were adult Thy1-GCaMP6f mice (Jackson Labs, strain code: 025393, C57BL/6J-Tg(Thy1-GCaMP6f)GP5.17Dkim/J)). The mice were implanted with bilateral cranial windows, one window on each of the hemispheres, and allowed to recover for two weeks. Following recovery, mice underwent a habituation training in a custom imaging cradle to get accustomed to the imaging setup and environment. Pre-stroke control measures were obtained one week prior to stroke and photothrombotic stroke was performed on Day0 of the experiment. Following photothrombosis, mice were imaged longitudinally at Day2, Week1, Week2, and Week4 to span both the acute and chronic phases of stroke recovery. To correlate the cortical measures to a behavior metric, forelimb asymmetry was measured using the cylinder test at each of the imaging time points. The timeline of experiments is outlined in Fig. 1a.

### Animal preparation

A bilateral cranial window exposing both hemispheres of the brain was implanted in all mice to determine the effect of stroke on both the ipsilesional and contralesional hemispheres. The surgical procedure for implantation of bilateral cranial windows followed a similar procedure to unilateral windows that has been previously described^69^. Briefly, mice were injected with Buprenorphine subcutaneously 1 hour prior to the start of surgery. During surgery, mice were anesthetized with isoflurane (3% at induction and 1-1.5% for maintenance with 1L/min oxygen) and body temperature was maintained at 37°C. Respiratory rate and toe pinch were used to monitor the depth of anesthesia throughout the surgical procedure. After incision of the scalp, a round aluminum head post, 12mm in diameter, was attached to the intact skull with dental acrylic. A craniotomy was the performed on one hemisphere of the brain in order to remove the skull. A half-skull-shaped curved glass (modified from Crystal Skull^70^, LabMaker, Germany) was used to cover the surface of the brain and then sealed with optical glue and dental acrylic. The craniotomy and glass procedure were repeated on the other hemisphere of the brain in order to create a bilateral cranial window implant. Recovery procedures were followed according to the guidelines provided by Boston University. After a two-week recovery period from surgery, mice were trained to remain head-fixed for up to 90 min for approximately 10 days. All experiments are done in awake head-fixed mice.

### Simultaneous hemodynamic and calcium imaging

To evaluate local and global changes in neurovascular coupling post-stroke simultaneous measures of hemodynamic and neural activity were obtained during forelimb sensory stimulation and resting state. The instrumentation, task setup, and data analysis pipeline for measuring cortical hemodynamics has been outlined previously^69^. Fig. 1a shows a simplified schematic of the imaging setup. Intrinsic optical signal imaging was used to assess changes to oxy and deoxy hemoglobin, HbO and HbR respectively, for the hemodynamic measure, and fluorescence GCaMP imaging was performed to assess changes in calcium dynamics as a measure of neural activity. The cortical windows were illuminated sequentially with 470 nm, 530 nm, and 625 nm LEDs (MXL3-C1, Thorlabs, X is the center wavelength), where the 470 nm LED was used for GCaMP excitation and the 530 nm and 625 nm LEDs were used for calculations of oxy and deoxy hemoglobin. A 500 nm long pass filter (FELH0500, Thorlabs) placed along the detection path blocked out any GCaMP excitation light. Images were collected by a sCMOS camera (Hamamatsu ORCA-Flash 4.0 V3) at 15 Hz, 5 Hz per wavelength, with an exposure time of 50 msec. For resting state, spontaneous activity was obtained for 8 min. For sensory stimulation, two imaging session were performed at each time point pre- and post-stroke, one where the contralateral (affected) forelimb was stimulated and the second where the ipsilateral (unaffected) forelimb was stimulated. Each stimulation session consisted of 20 trials where each trial was obtained in a block-design fashion and consisted of 5 seconds of baseline, followed by 5 seconds of 3Hz air-puff stimulation, followed by 20 seconds of recovery. A custom MATLAB code was used to synchronize and trigger the sequential LEDs, camera acquisition, and air puff stimulation. Raw images at 530 nm and 625 nm were analyzed for changes in oxy- and deoxy-hemoglobin using the modified Beer-Lambert relationship as described previously^69,71^. Calcium dynamics were analyzed as a change in fluorescence over time from the interspersed raw images excited at 470 nm. The fluorescence data were corrected for hemodynamic crosstalk as hemodynamic changes contaminate the fluorescence signal and both the excitation and emission wavelengths. The correction algorithm used has been previously described and modified from Ma et al^32^. The correction implemented estimates the attenuation experienced by the GCaMP signal from the simultaneously obtained changes in oxy and deoxy hemoglobin concentration. The change in calcium concentration is approximately equal to the change in GCaMP fluorescence scaled by a time-varying hemoglobin absorption factor at both the GCaMP excitation and emission wavelengths. The pathlength factor used for correction is obtained from Monte Carlo simulations of photon transport using the Monte Carlo eXtreme (MCX) platform^72,73^. The absorption and scattering coefficients used for the MCX simulation were obtained from spatial frequency domain imaging (described below). For pre-stroke imaging, a single absorption and scattering coefficient, yielding a single pathlength, was used for correction of all pixels. After stroke, the absorption and scattering coefficients used were determined on a semi pixel-by-pixel basis. This modified correction technique was introduced in order to account for changes in tissue optical properties after stroke^74,75^. A Monte Carlo simulation was run on any pixel that had a scattering coefficient that was 30% larger than the mean scattering coefficient of the control animals, using the respective absorption and scattering coefficients of that pixel. This new pathlength was used for the correction of pixels within the stroke region that had increased scattering. The attenuation correction applied spatial maps and temporal traces are shown in Fig. 1b.

### Spatial frequency domain imaging

To capture the spatial extent of the stroke core longitudinally as well as to aid in fluorescence correction for hemodynamic crosstalk, SFDI was performed pre-stroke and at each time point post-stroke. The instrumentation, acquisition, and analysis to obtain absorption and scattering coefficients of the tissue have been described previously^74^. Spatially varying sinusoidal patterns were projected onto the cranial window by a digital micromirror device (DMD), and the reflected light was imaged by the sCMOS camera. Two spatial frequencies (0 and 0.4 mm^-1^) were projected at three phases (0, 120, and 240 deg). The acquired images were processed offline using MATLAB. The intensity at each spatial frequency was demodulated and calibrated to a reference phantom to obtain the diffuse reflectance. A two-frequency lookup table was generated by Monte Carlo simulations at the two frequencies used for imaging from which absorption and scattering coefficients were extracted. To obtain the spatial extent of the stroke core, the relative change in scattering coefficient post-stroke was calculated with respect to pre-stroke scattering, and a semi-automatic contour was applied using a custom MATLAB code, to create a stroke core outline. This core outline was used as the boundary for the start of the peri-infarct zone^74^. The peri-infarct zone was defined as the region that extended 0.5mm outward from the stroke core outline. SFDI was also used in the correction of GCaMP for hemodynamic crosstalk. The absorption and scattering properties obtained at each time point post-stroke were used to run the Monte Carlo simulation to determine the pathlength of light travelled in tissue. This pathlength is then used in the correction algorithm to scale the GCaMP signal, based on the time-varying changes in hemodynamic absorption, for accurate estimation of calcium dynamics.

### Resting state functional connectivity analysis

Global network connectivity changes following stroke were assessed using resting state functional connectivity as described previously by a number of groups^40,45,47^. Time traces of HbO and GCaMP were bandpass filtered into two frequency bands, the typically used infraslow (0.008-0.09 Hz) frequency band and a higher frequency band (0.09-0.4 Hz) and regressed to remove any global fluctuations in the signal. To evaluate the strength of network connections to the affected forelimb region, a seed was placed in the center of the original forelimb somatosensory region of the affected hemisphere. The seed time trace was calculated by averaging the time trace within 0.25 mm of the seed location and connectivity was assessed by calculating the correlation between the seed time trace and the time course of every other pixel. By averaging the positive correlation coefficients between the forelimb seed and all pixels that lie in the contralesional forelimb region we calculated a forelimb connectivity map^47^. Interhemispheric connectivity maps were calculated by correlating each pixel within the affected hemisphere with its mirror pixel, mirrored along the midline, in the unaffected hemisphere. The interhemispheric connectivity index was then calculated by averaging all the pixels within the homotopic map of the affected hemisphere^47^. To assess the overall connectivity of the brain, global connectivity maps were generated by calculating the correlation of each pixel with every other pixel and then assigning the average positive correlation coefficient to that pixel. From the global connectivity maps, a global connectivity index was calculated by taking the mean of the correlation coefficients for all pixels within the map^47^.

### Neurovascular coupling

To assess the relationship between neural activity and hemodynamics, neurovascular coupling was modeled using linear least-squares deconvolution^32^. The cortical hemodynamic response is known to be a linear convolution of the cortical neural activity and an impulse response function (IRF). The impulse response function, also called the hemodynamic response function, is the hemodynamic response to a neural stimulus. In a linear system, the convolution can be expressed as *Y* = *X*\**h*, and can be represented as:

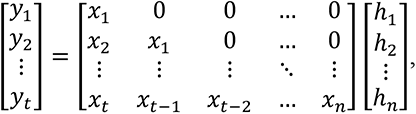

where *X* is the input to the system, which is the corrected GCaMP fluorescence signal, and the length *n* used is 15 sec (from -5 sec to 10 sec), *y* is the output of the system, which is the hemodynamic signal, and *h* is the system’s impulse function. A direct solution to the linear system could result in an ill-conditioned matrix and therefore a regularization term is added and the solution is obtained by minimizing the cost function and setting the derivative of the cost function to zero, as described previously, and is given by:

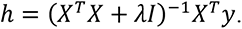

The regularization term λ was chosen to be 0.1 through all the analysis. The deconvolution was performed on a pixel-by-pixel basis at each time point post-stroke.

### Targeted photothrombosis

Focal cerebral ischemia was performed using an optimized photothrombosis method described previously^69^. A distal branch of the middle cerebral artery supplying the forelimb somatosensory region, determined through pre-stroke forelimb stimulation, was targeted for occlusion. A 520nm laser diode with axial and lateral parameters of 104 µm and 6 µm was tuned to a minimal post-objective power of 0.6 mW. These parameters were designed to occlude only the target vessel and prevent laser damage to the surrounding tissue, thus ensuring that the ischemia procedure was physiological in nature. Real-time changes to cerebral blood flow (CBF) were monitored through laser speckle contrast imaging (LSCI). Ten minutes of baseline CBF was obtained following which the mouse was lightly anesthetized to inject Rose Bengal (100 µl, 15 mg/ml in saline) retroorbitally. The mouse was then immediately taken off isoflurane and allowed to recover, which was determined by a return of CBF to baseline and the mouse exhibiting natural behaviors such as whisking. Following recovery the green laser was turned on until the target vessel was occluded, as indicated by the target branch disappearing on LSCI. Once the target branch was occluded, the laser power was reduced to 0.5 mW for an additional minute and then turned off. If at any point the target branch started flowing again, the laser was turned back on until occlusion. Additionally, as described previously, two collateral branches were also targeted to obtain a stable infarct. The procedure was followed for 1 hour from the initiation of photothrombosis.

### Behavioral testing

The cylinder test was used in all mice to assess behavioral deficit in forelimb use over the course of 4 weeks following stroke. Two sessions of pre-stroke testing was obtained the week before stroke induction to assess basal preference in forepaw use. Following photothrombotic stroke, mice were tested at 2 days, 1 week, 2 weeks, and 4 weeks. Each testing session involved placing a mouse in a clear glass cylinder and videotaping its natural behavior from below for 15 minutes. Forelimb use was assessed by counting the number of times the mouse used each forelimb to make first contact with the cylinder wall during rears. Asymmetry in forelimb use after stroke was quantified as a percent change from baseline use of the contralateral (affected) forelimb. Change from baseline was used to compensate for the fact that some mice have a preference for one paw over the other even before a stroke.

### Data analysis and Statistics

All data was analyzed offline using custom MATLAB codes. Image analysis for SFDI, calcium fluorescence, and evoked and resting-state intrinsic optical signal imaging has been outlined in previous sections. The dice similarity coefficient for area overlap in evoked responses and RSFC is calculated using the matlab function dice.m. The dice coefficient is twice the ratio of the intersection of two binary images and the sum of the number of elements in each image, given by:

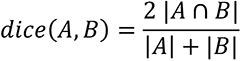

Goodness-of-fit correlation and significance for stimulus evoked response magnitudes of GCaMP and hemodynamics were made using a linear fit. All statistical analyses were made using MATLAB with *post hoc* comparisons using t-tests. A two sample students t-test was performed for comparing data points with pre-stroke data (matlab function: ttest2).

## Data and code availability statement

The datasets generated and/or analyzed during this study and corresponding code that support the findings of this study are available from the corresponding author upon request.

## Acknowledgments

This work was supported by the National Institute of Health [R01-EB021018, R01-NS108472, R01-MH111359].

## Author contributions

Conceptualization: SS, KK, EE, DAB

Methodology: SS, JJ, SK, KK, EE, DAB

Investigation: SS, ShS

Visualization: SS, DAB

Supervision: DAB, AD, CA

Writing: SS, EE, CA, AD, DAB

---

**Supplementary figure 1:**
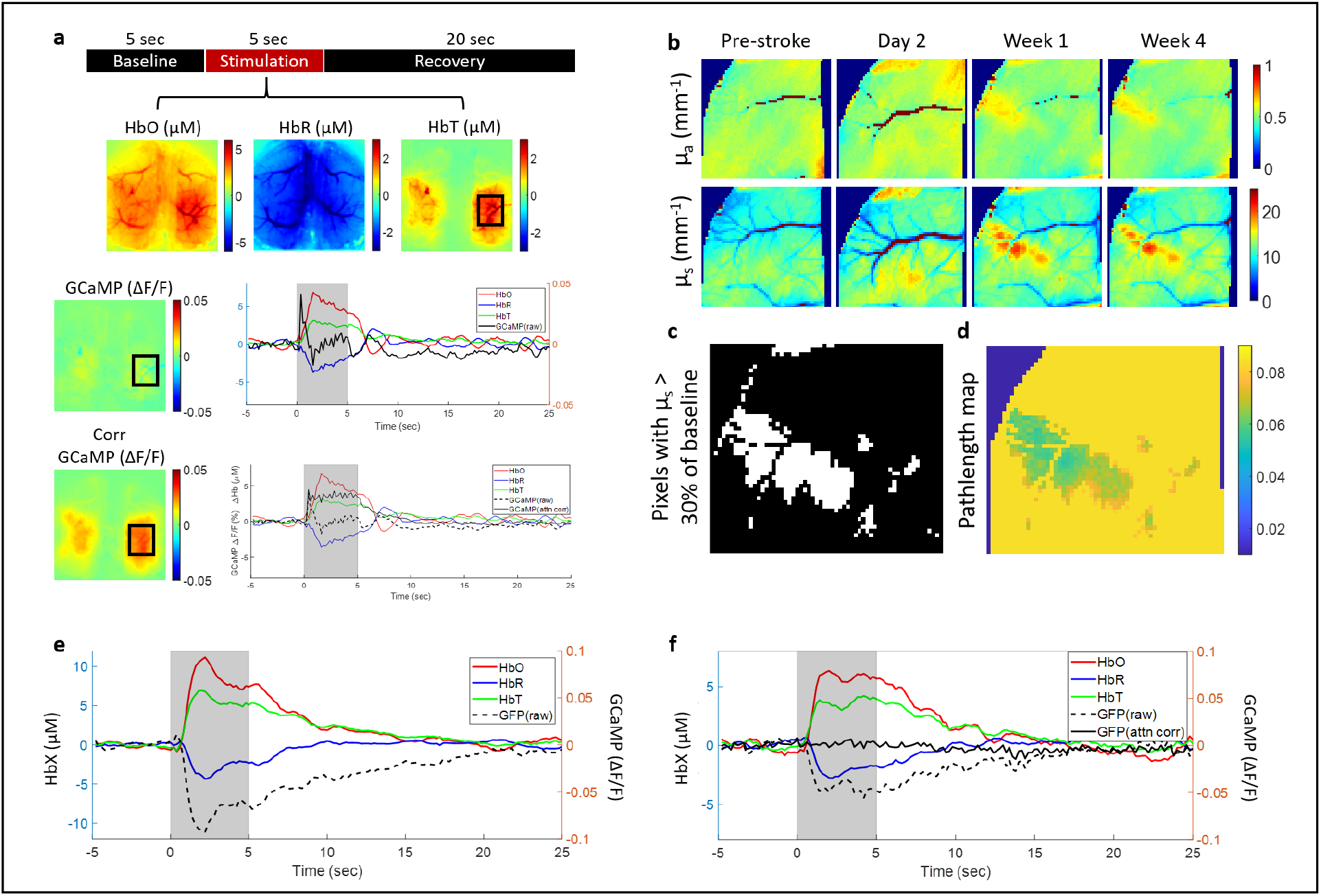
Fluorescence correction for hemodynamic crosstalk. (a) Top: Block design of single sensory stimulation trial and spatial hemodynamic response maps for HbO, HbR, and HbT. Middle: Raw GCaMP response map during 5 sec of sensory stimulation and time course of trial averaged data for GCaMP and hemodynamics from ROI marked in black box. Uncorrected GCaMP shows rise in fluorescence at the start of stimulation but begins to decrease with the rise of hemodynamic response. Bottom: Spatial map of GCaMP corrected for hemodynamic crosstalk. Note the appearance of response compared to uncorrected GCaMP in spatial map. Time course of corrected GCaMP overlaid with uncorrected GCaMP and hemodynamics. Note that GCaMP is now elevated for the full stimulation period. (b) Absorption and scattering coefficients obtained from SFDI before and after stroke and used in the correction algorithm in the form of pathlength factor. Stroke leads to increases in the scattering signal that needs to be accounted for accurate correction due to its effect on pathlength. (c) Binary maps of all pixels that have scattering coefficient greater than 30% of baseline scattering. The scattering and absorption coefficients from these pixels are used in the Monte Carlo simulation to obtain pathlength. (d) Spatial map of pathlength factors obtained from Monte Carlo simulations and used in the correction algorithm. (e,f) Validation of correction algorithm with cellular fluorescent marker GFP. (e) GFP signal overlaid with hemodynamics during 5sec of sensory stimulation. GFP drops in association with hemodynamic increase. (f) Correction applied to GFP signal during sensory stimulation. Corrected GFP is a flat line as expected since GFP fluorescence is not altered with neural activity or hemodynamics.

**Supplementary figure 2:**
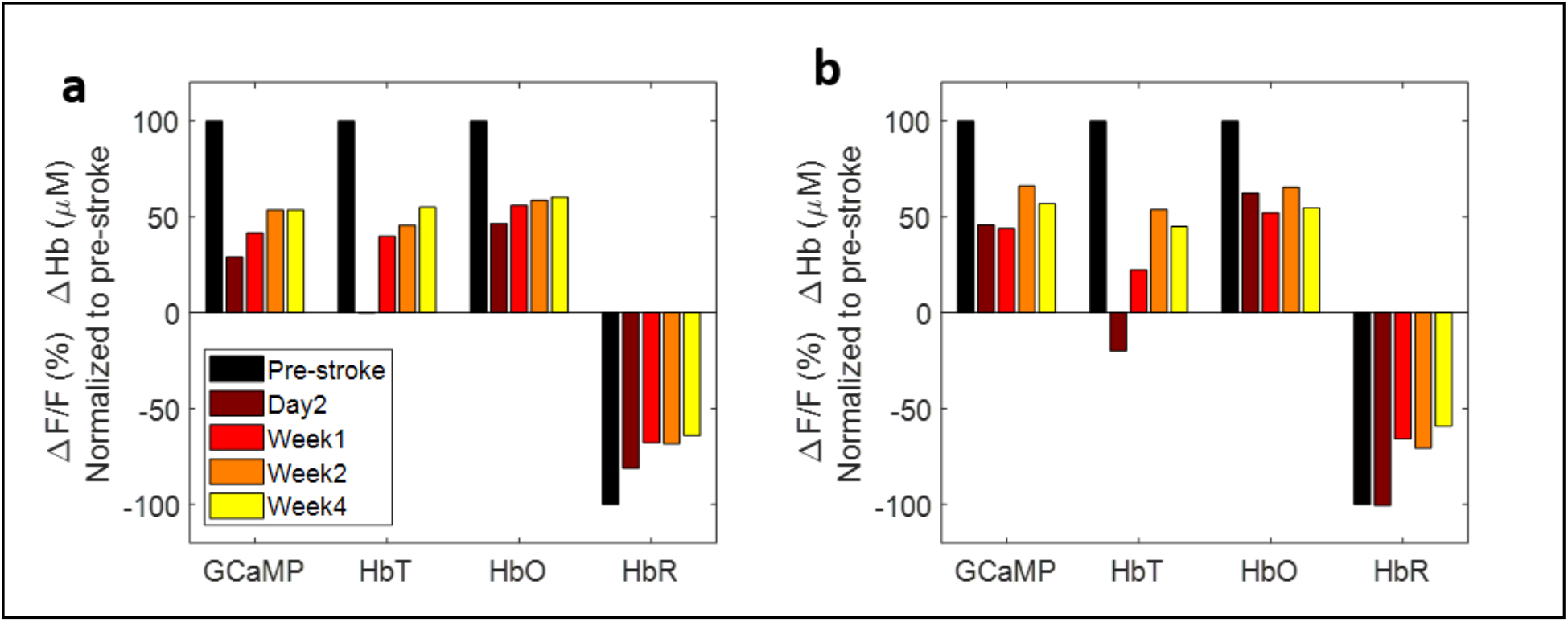
Responses in the affected hemisphere normalized to pre-stroke during stimulation of the impaired (a) and unimpaired (b) forelimb. HbT response is more sensitive to the stroke compared with HbO and HbR.

**Supplementary figure 3:**
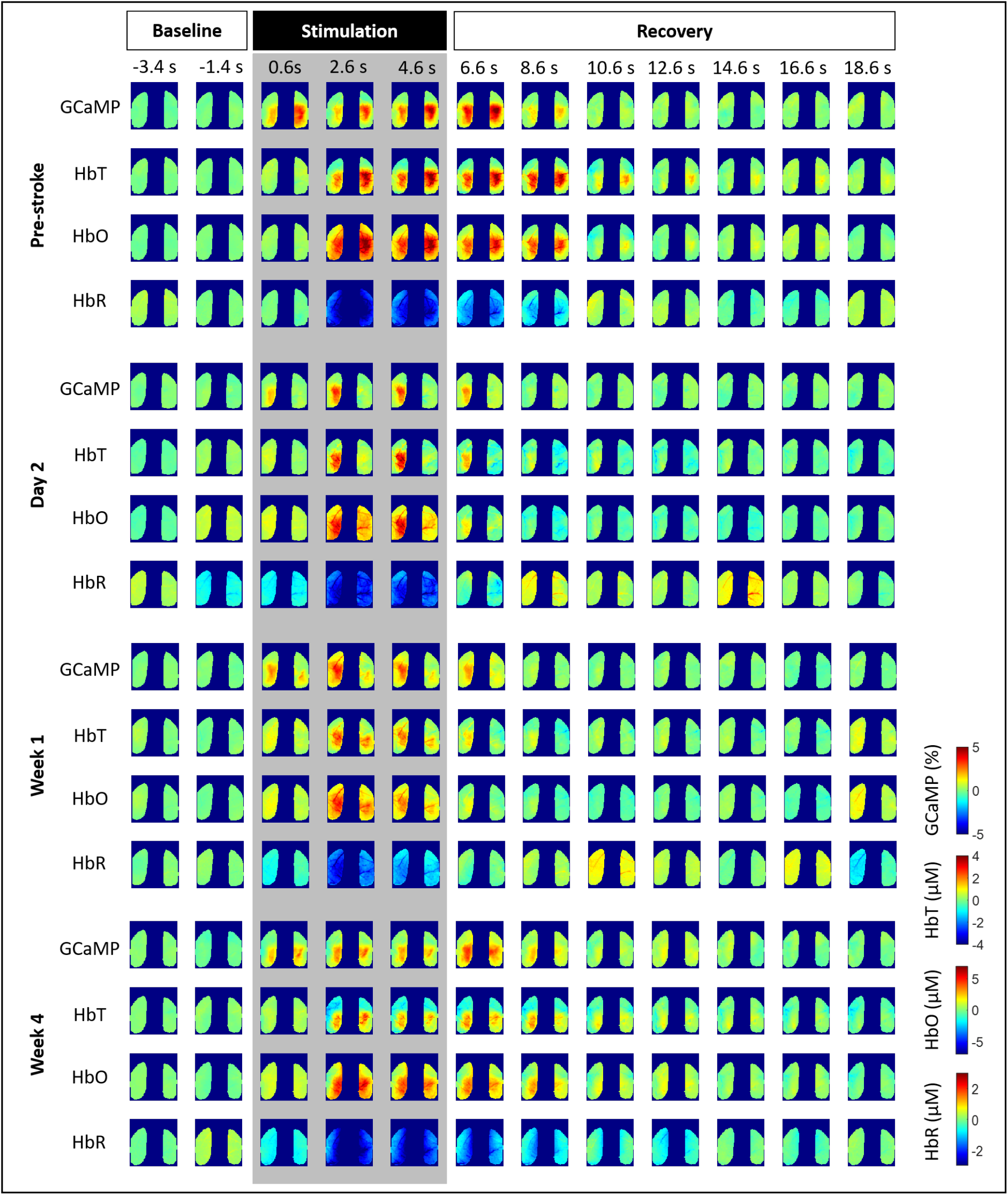
Spatial maps of GCaMP and hemodynamic responses over time during sensory stimulation.

**Supplementary figure 4:**
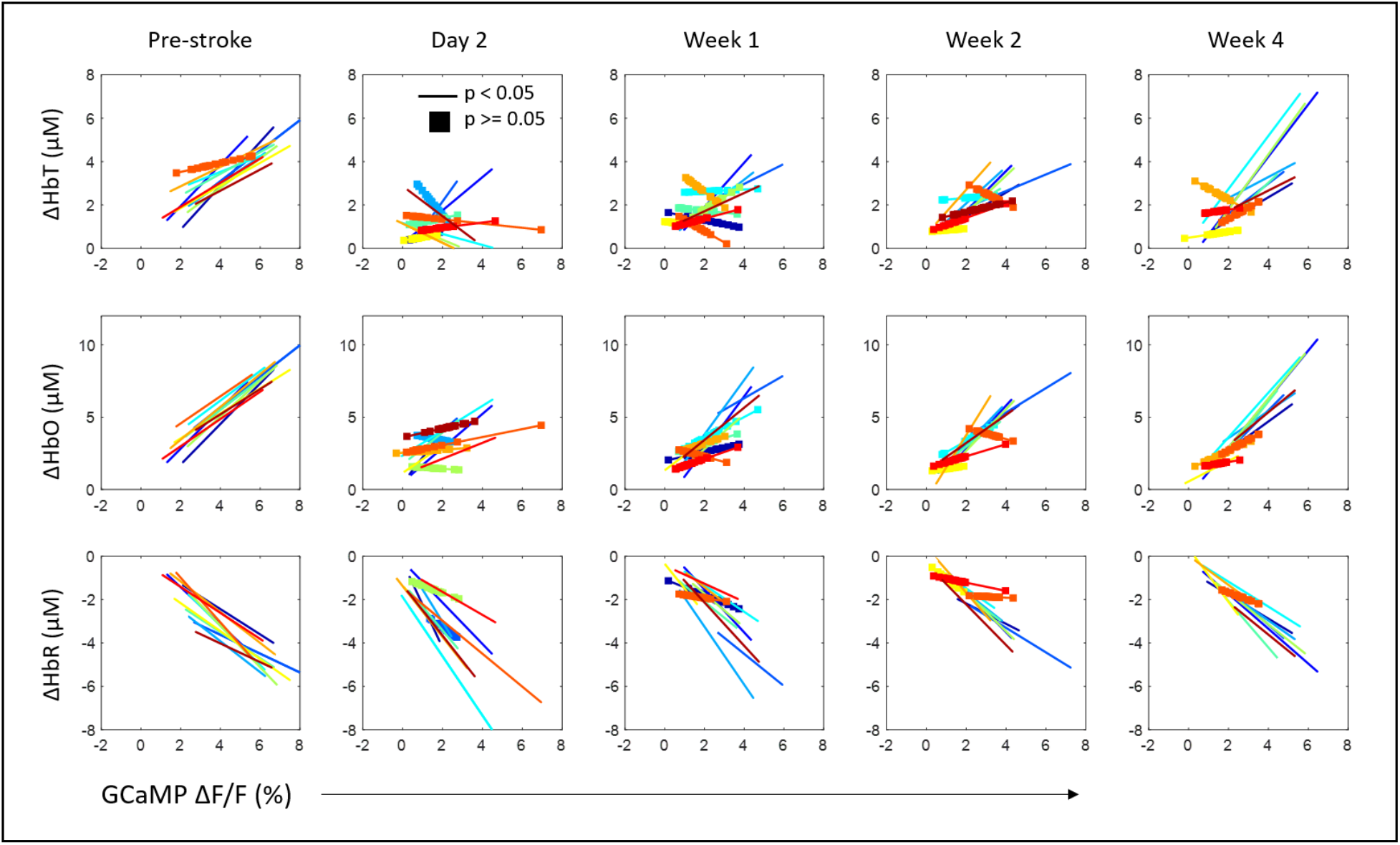
Correlation of calcium and hemodynamic evoked responses to sensory stimulation of the affected forelimb color-coded by mouse. Mice with significant correlation in response magnitudes of calcium and hemodynamics are shown as solid lines and mice whose responses were not correlated are shown with filled squares. Note that the animals that did not show correlation at week 4 after stroke also lacked correlation in the acute phase of stroke at week 1.

**Supplementary figure 5:**
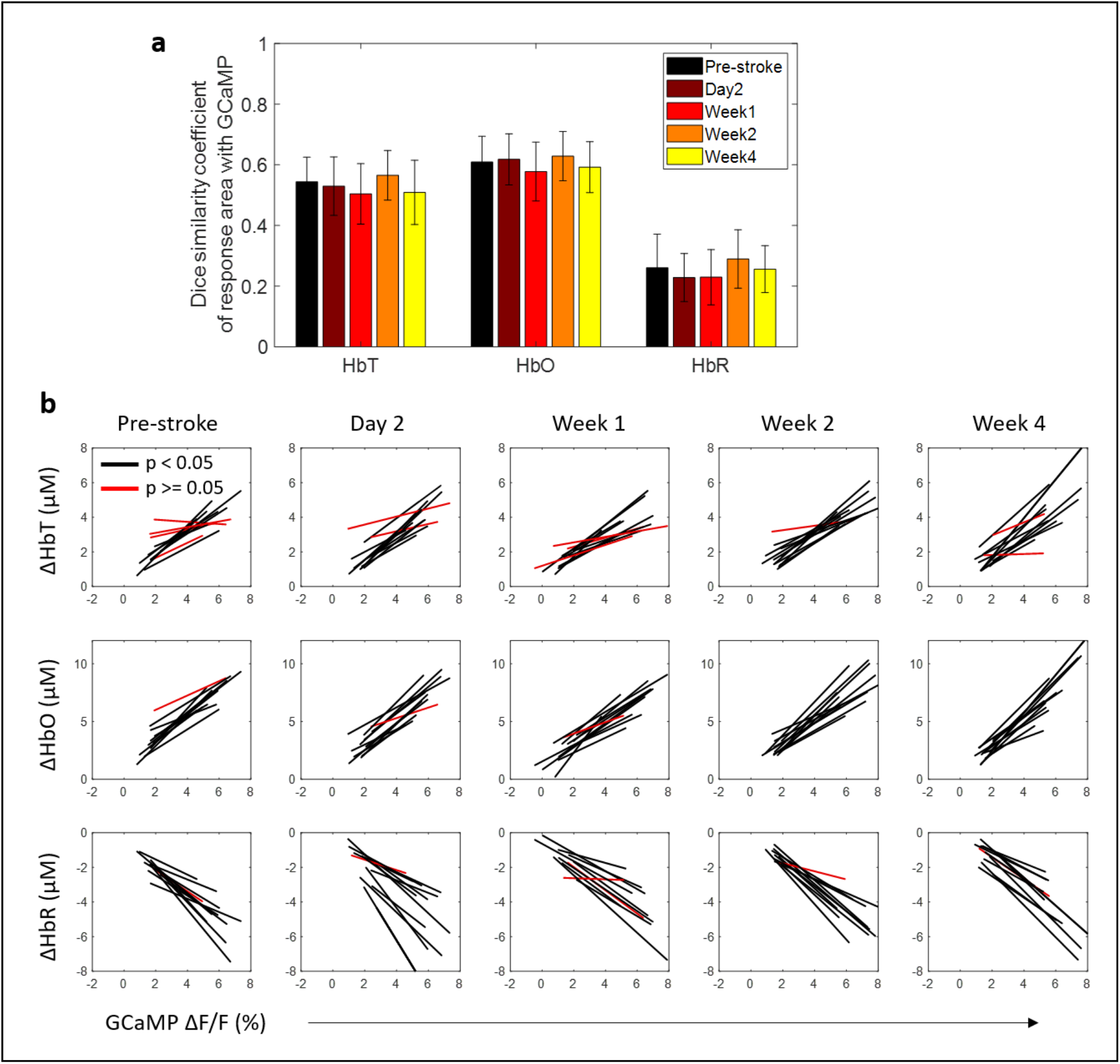
Responses within the unaffected hemisphere during stimulation of the unaffected forelimb. (a) Dice similarity coefficient between GCaMP response areas with each hemodynamic measure. There was no change in similarity of response area after stroke. (b) Correlation of calcium and hemodynamic evoked responses in the unaffected forelimb to sensory stimulation of the unaffected forelimb.

**Supplementary figure 6:**
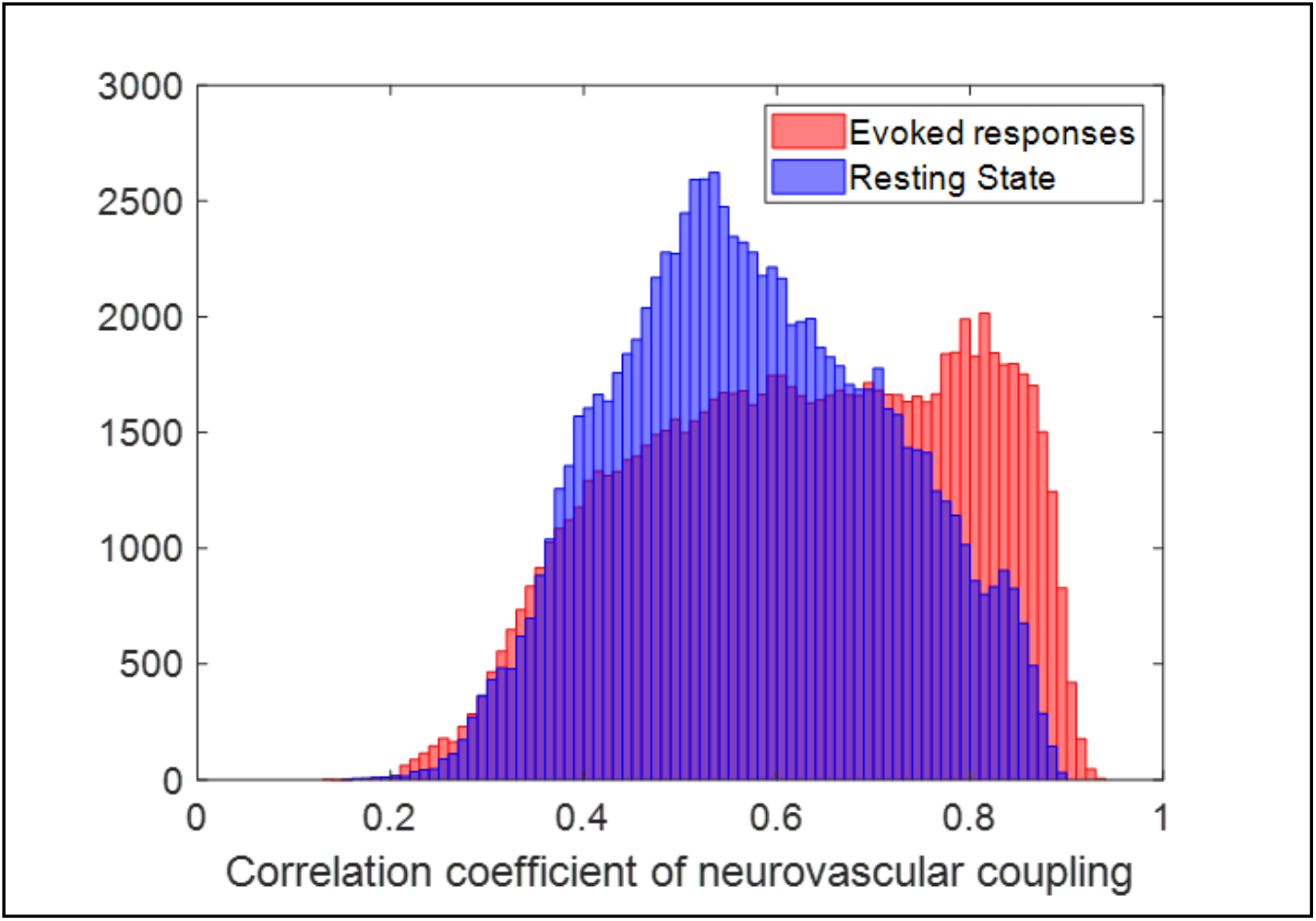
Pearson’s correlation coefficient of neurovascular coupling in healthy pre-stroke animals during sessions with evoked responses and resting-state sessions.

**Supplementary figure 7:**
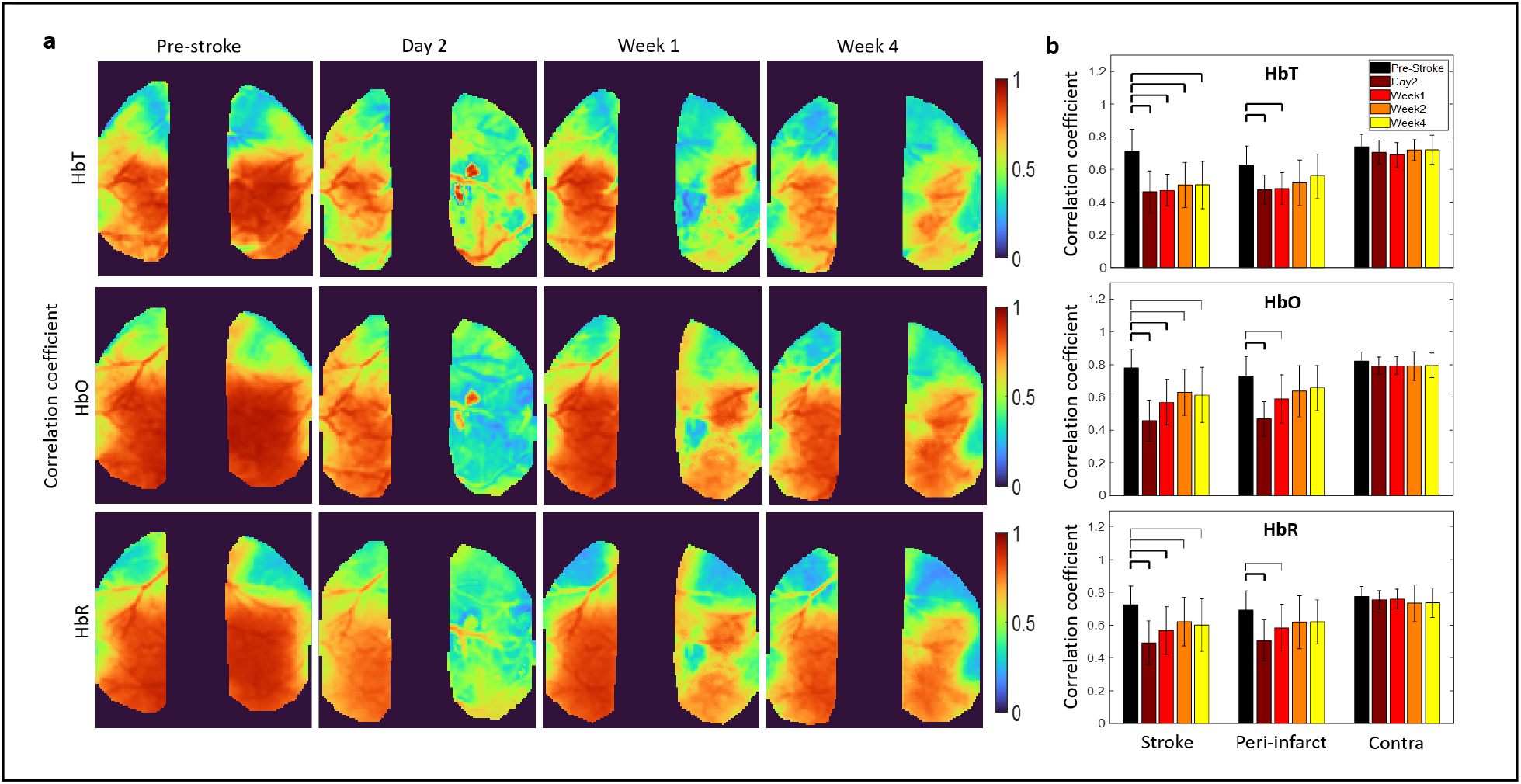
(a) Pixel-by-pixel Pearson’s correlation coefficient between measured and predicted HbT (top), HbO (middle), and HbR (bottom). Predicted HbX is obtained by convolving the GCaMP signal at each time point with the HRF obtained for that specific time point and pixel. (h) Pearson’s correlation coefficient quantified across all mice within the stroke core, peri-infarct, and contralesional forelimb region. Thick bars: p<0.01, thin bars: p<0.05.

**Supplementary figure 8:**
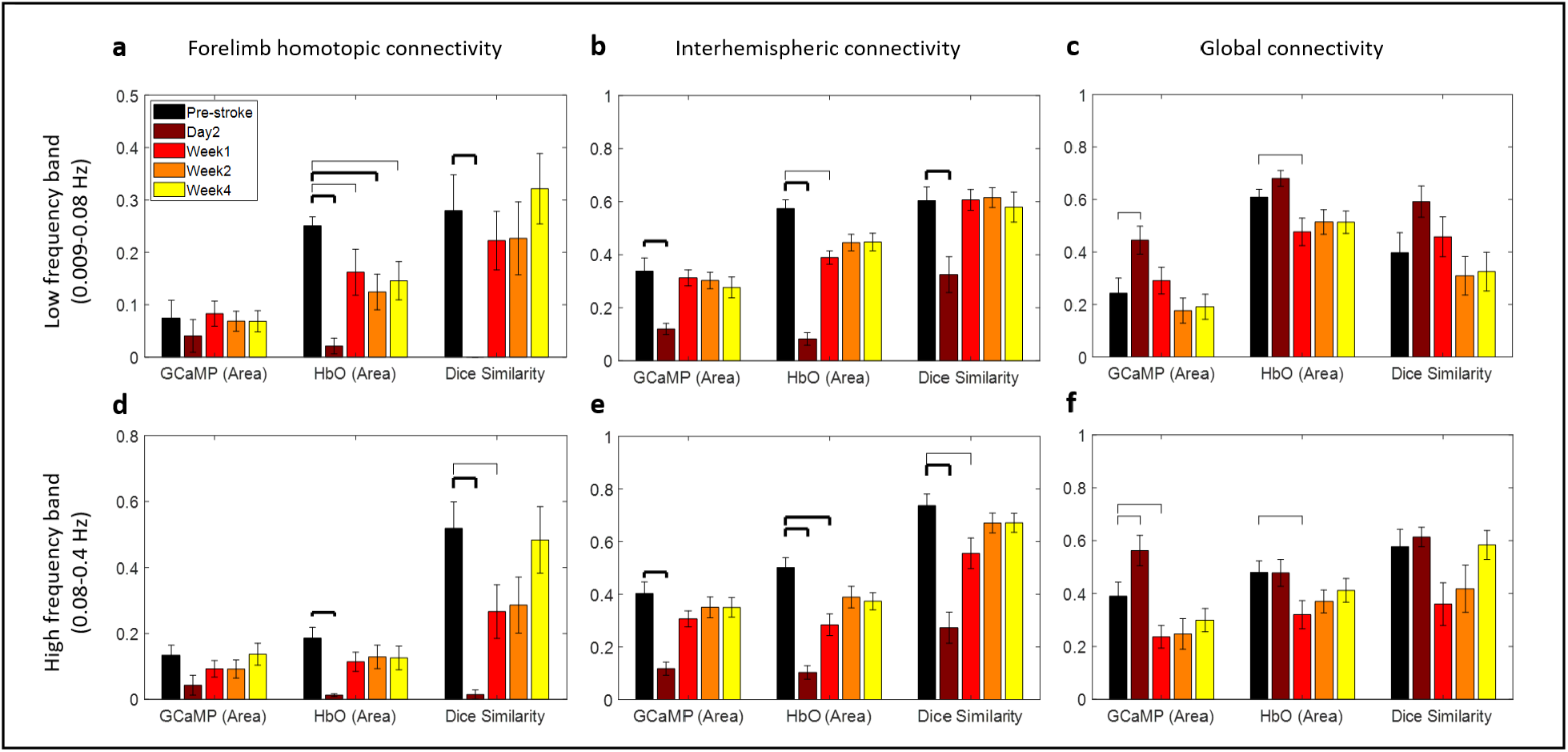
RSFC proportional area and dice coefficient analysis at threshold of 0.4. Each figure shows the proportional area of GCaMP and HbO above the correlation coefficient equal to 0.4 and the dice similarity between the GCaMP and HbO at 0.4. (a) Forelimb homotopic connectivity in the low frequency band and (d) high frequency band. (b) Interhemispheric connectivity in the low frequency band and (e) high frequency band. (c) Global connectivity in the low frequency band and (f) high frequency band.

## References

1. Moskowitz, M. A., Lo, E. H. & Iadecola, C. The science of stroke: Mechanisms in search of treatments. Neuron 67, 181–198 (2010).

2. Lo, E. H., Moskowitz, M. A. & Jacobs, T. P. Exciting, radical, suicidal: How brain cells die after stroke. Stroke 36, 189–192 (2005).

3. Sharma, N. & Cohen, L. G. Recovery of motor function after stroke. Dev. Psychobiol. 54, 254–262 (2012).

4. Cramer, S. C. Repairing the human brain after stroke: I. Mechanisms of spontaneous recovery. Ann. Neurol. 63, 272–287 (2008).

5. Cassidy, J. M. & Cramer, S. C. Spontaneous & Therapeutic-Induced Mechanisms of Functional Recovery After Stroke. Transl. Stroke Res. 8, 33–46 (2017).

6. Jones, T. A. Motor compensation and its effects on neural reorganization after stroke. Nat. Rev. Neurosci. 18, 267–280 (2017).

7. Moseley, M. Mri of stroke. Imaging 41, 410–414 (2010).

8. Veldsman, M., Cumming, T. & Brodtmann, A. Beyond BOLD: Optimizing functional imaging in stroke populations. Hum. Brain Mapp. 36, 1620–1636 (2015).

9. Lake, E. M. R., Bazzigaluppi, P. & Stefanovic, B. Functional magnetic resonance imaging in chronic ischaemic stroke. Philos. Trans. R. Soc. B Biol. Sci. 371, 1–11 (2016).

10. Johansen-Berg, H. et al. Correlation between motor improvements and altered fMRI activity after rehabilitative therapy. Brain 125, 2731–2742 (2002).

11. Pineiro, R., Pendlebury, S., Johansen-Berg, H. & Matthews, P. M. Altered hemodynamic responses in patients after subcortical stroke measured by functional MRI. Stroke 33, 103– 109 (2002).

12. Carter, A. R. et al. Resting state inter-hemispheric fMRI connectivity predicts performance after stroke. Ann. Neurol. 67, NA-NA (2009).

13. Corbetta, M. Functional connectivity and neurological recovery. Dev. Psychobiol. 54, 239–253 (2012).

14. Kleinfeld, D. et al. A guide to delineate the logic of neurovascular signaling in the brain. Front. Neuroenergetics 3, 1–9 (2011).

15. Buxton, R. B., Griffeth, V. E. M., Simon, A. B. & Moradi, F. Variability of the coupling of blood flow and oxygen metabolism responses in the brain: A problem for interpreting BOLD studies but potentially a new window on the underlying neural activity. Front. Neurosci. 8, 1–6 (2014).

16. Logothetis, N. K., Pauls, J., Augath, M., Trinath, T. & Oeltermann, A. Neurophysiological investigation of the basis of the fMRI signal. Nature 412, 150–157 (2001).

17. Kunz, A. & Iadecola, C. Cerebral vascular dysregulation in the ischemic brain. Handb. Clin. Neurol. 92, 283–305 (2009).

18. Girouard, H. & Iadecola, C. Neurovascular coupling in the normal brain and in hypertension, stroke, and Alzheimer disease Regulation of the Cerebral Circulation stroke, and Alzheimer disease. 10021, 328–335 (2012).

19. Weber, R. et al. Early prediction of functional recovery after experimental stroke: Functional magnetic resonance imaging, electrophysiology, and behavioral testing in rats. J. Neurosci. 28, 1022–1029 (2008).

20. Shih, Y. Y. I. et al. Imaging neurovascular function and functional recovery after stroke in the rat striatum using forepaw stimulation. J. Cereb. Blood Flow Metab. 34, 1483–1492 (2014).

21. Carmichael, S. T. Rodent models of focal stroke: Size, mechanism, and purpose. NeuroRX 2, 396–409 (2005).

22. Bacigaluppi, M., Comi, G. & Hermann, D. M. Animal models of ischemic stroke. Part two: modeling cerebral ischemia. Open Neurol. J. 4, 34–38 (2010).

23. Sommer, C. J. Ischemic stroke: experimental models and reality. Acta Neuropathol. 133, 245–261 (2017).

24. Winship, I. R. & Murphy, T. H. In Vivo Calcium Imaging Reveals Functional Rewiring of Single Somatosensory Neurons after Stroke. J. Neurosci. 28, 6592–6606 (2008).

25. Brown, C. E., Aminoltejari, K., Erb, H., Winship, I. R. & Murphy, T. H. In Vivo Voltage-Sensitive Dye Imaging in Adult Mice Reveals That Somatosensory Maps Lost to Stroke Are Replaced over Weeks by New Structural and Functional Circuits with Prolonged Modes of Activation within Both the Peri-Infarct Zone and Distant Sites. J. Neurosci. 29, 1719–1734 (2009).

26. Harrison, T. C., Silasi, G., Boyd, J. D. & Murphy, T. H. Displacement of sensory maps and disorganization of motor cortex after targeted stroke in mice. Stroke 44, 2300–2306 (2013).

27. Clarkson, A. N. et al. Multimodal examination of structural and functional remapping in the mouse photothrombotic stroke model. J. Cereb. Blood Flow Metab. 33, 716–723 (2013).

28. Lim, D. H., LeDue, J. M., Mohajerani, M. H. & Murphy, T. H. Optogenetic Mapping after Stroke Reveals Network-Wide Scaling of Functional Connections and Heterogeneous Recovery of the Peri-Infarct. J. Neurosci. 34, 16455–16466 (2014).

29. Bauer, A. Q. et al. Optical imaging of disrupted functional connectivity following ischemic stroke in mice. Neuroimage 99, 388–401 (2014).

30. Park, C. H. et al. Longitudinal changes of resting-state functional connectivity during motor recovery after stroke. Stroke 42, 1357–1362 (2011).

31. Schrandt, C. J., Kazmi, S. S., Jones, T. A. & Dunn, A. K. Chronic Monitoring of Vascular Progression after Ischemic Stroke Using Multiexposure Speckle Imaging and Two-Photon Fluorescence Microscopy. J. Cereb. Blood Flow Metab. 35, 933–942 (2015).

32. Ma, Y. et al. Resting-state hemodynamics are spatiotemporally coupled to synchronized and symmetric neural activity in excitatory neurons. Proc. Natl. Acad. Sci. 113, E8463–E8471 (2016).

33. Wright, P. W. et al. Functional connectivity structure of cortical calcium dynamics in anesthetized and awake mice. PLoS One 12, 1–27 (2017).

34. Tian, L. et al. Imaging neural activity in worms, flies and mice with improved GCaMP calcium indicators. Nat. Methods 6, 875–881 (2009).

35. Dana, H. et al. Thy1-GCaMP6 transgenic mice for neuronal population imaging in vivo. PLoS One 9, (2014).

36. Ma, Y. et al. High-speed, wide-field optical mapping (WFOM) of neural activity and brain haemodynamics: Considerations and novel approaches. Under Rev. 371, 20150360 (2016).

37. Montgomery, M. K. et al. Glioma-Induced Alterations in Neuronal Activity and Neurovascular Coupling during Disease Progression. Cell Rep. 31, 107500 (2020).

38. Lake, E. M. R. et al. Simultaneous cortex-wide fluorescence Ca2+ imaging and whole-brain fMRI. Nat. Methods 17, 1262–1271 (2020).

39. Winder, A. T., Echagarruga, C., Zhang, Q. & Drew, P. J. Weak correlations between hemodynamic signals and ongoing neural activity during the resting state. Nat. Neurosci. 20, 1761–1769 (2017).

40. Wright, P. W. et al. Functional connectivity structure of cortical calcium dynamics in anesthetized and awake mice. PLoS One 12, (2017).

41. Laaksonen, K. et al. Alterations in Spontaneous Brain Oscillations during Stroke Recovery. PLoS One 8, (2013).

42. Goltsov, A. et al. Bifurcation in blood oscillatory rhythms for patients with ischemic stroke: A small scale clinical trial using laser Doppler flowmetry and computational modeling of vasomotion. Front. Physiol. 8, 1–11 (2017).

43. Grefkes, C. & Fink, G. R. Reorganization of cerebral networks after stroke: New insights from neuroimaging with connectivity approaches. Brain 134, 1264–1276 (2011).

44. Grefkes, C. & Fink, G. R. Connectivity-based approaches in stroke and recovery of function. Lancet Neurol. 13, 206–216 (2014).

45. Chung, D. Y. et al. Subarachnoid hemorrhage leads to early and persistent functional connectivity and behavioral changes in mice. J. Cereb. Blood Flow Metab. 41, 975–985 (2021).

46. Kura, S. et al. Intrinsic optical signal imaging of the blood volume changes is sufficient for mapping the resting state functional connectivity in the rodent cortex. J. Neural Eng. c, (2018).

47. Xie, H. et al. Differential effects of anesthetics on resting state functional connectivity in the mouse. J. Cereb. Blood Flow Metab. 40, 875–884 (2020).

48. Angels Font, M., Arboix, A. & Krupinski, J. Angiogenesis, Neurogenesis and Neuroplasticity in Ischemic Stroke. Curr. Cardiol. Rev. 6, 238–244 (2010).

49. Boyd, L. A. et al. Biomarkers of stroke recovery: Consensus-based core recommendations from the Stroke Recovery and Rehabilitation Roundtable. Int. J. Stroke 12, 480–493 (2017).

50. Stinear, C. M. Prediction of motor recovery after stroke: advances in biomarkers. Lancet Neurol. 16, 826–836 (2017).

51. Rabiller, G., He, J. W., Nishijima, Y., Wong, A. & Liu, J. Perturbation of brain oscillations after ischemic stroke: A potential biomarker for post-stroke function and therapy. Int. J. Mol. Sci. 16, 25605–25640 (2015).

52. Huang, M. X. et al. Marked Increases in Resting-State MEG Gamma-Band Activity in Combat-Related Mild Traumatic Brain Injury. Cereb. Cortex 30, 283–295 (2020).

53. Intaglietta, M. Arteriolar Vasomotion: Implications for Tissue Ischemia. J. Vasc. Res. 28(suppl 1, 1–7 (1991).

54. Intaglietta, M. Vasomotion and flowmotion: physiological mechanisms and clinical evidence. Vasc. Med. Rev. **vmr**-1, 101–112 (1990).

55. Carmichael, S. T. Brain excitability in stroke: The yin and yang of stroke progression. Arch. Neurol. 69, 161–167 (2012).

56. Joy, M. T. & Carmichael, S. T. Encouraging an excitable brain state: mechanisms of brain repair in stroke. Nat. Rev. Neurosci. 22, 38–53 (2021).

57. Bütefisch, C. M., Netz, J., Weßling, M., Seitz, R. J. & Hömberg, V. Remote changes in cortical excitability after stroke. Brain 126, 470–481 (2003).

58. Mohajerani, M. H., Aminoltejari, K. & Murphy, T. H. Targeted mini-strokes produce changes in interhemispheric sensory signal processing that are indicative of disinhibition within minutes. Proc. Natl. Acad. Sci. 108, E183–E191 (2011).

59. Balbi, M. et al. Gamma frequency activation of inhibitory neurons in the acute phase after stroke attenuates vascular and behavioral dysfunction. Cell Rep. 34, 108696 (2021).

60. Adaikkan, C. et al. Gamma Entrainment Binds Higher-Order Brain Regions and Offers Neuroprotection. Neuron 102, 929–943.e8 (2019).

61. Rehme, A. K., Eickhoff, S. B., Wang, L. E., Fink, G. R. & Grefkes, C. Dynamic causal modeling of cortical activity from the acute to the chronic stage after stroke. Neuroimage 55, 1147–1158 (2011).

62. Rehme, A. K., Eickhoff, S. B., Rottschy, C., Fink, G. R. & Grefkes, C. Activation likelihood estimation meta-analysis of motor-related neural activity after stroke. Neuroimage 59, 2771–2782 (2012).

63. Carmichael, S. T. & Chesselet, M. F. Synchronous neuronal activity is a signal for axonal sprouting after cortical lesions in the adult. J. Neurosci. 22, 6062–6070 (2002).

64. White, B. R. et al. Imaging of functional connectivity in the mouse brain. PLoS One 6, (2011).

65. Isaacson, J. S. & Scanziani, M. How inhibition shapes cortical activity. Neuron 72, 231– 243 (2011).

66. Hamel, E. Perivascular nerves and the regulation of cerebrovascular tone. J. Appl. Physiol. 100, 1059–1064 (2006).

67. Attwell, D. et al. Glial and neuronal control of brain blood flow. Nature 468, 232–243 (2010).

68. Lo, E. H., Dalkara, T. & Moskowitz, M. A. Neurological diseases: Mechanisms, challenges and opportunities in stroke. Nat. Rev. Neurosci. 4, 399–414 (2003).

69. Sunil, S. et al. Awake chronic mouse model of targeted pial vessel occlusion via photothrombosis. Neurophotonics 7, 1–18 (2020).

70. Kim, T. H. et al. Long-Term Optical Access to an Estimated One Million Neurons in the Live Mouse Cortex. Cell Rep. 17, 3385–3394 (2016).

71. Dunn, A. K., Devor, A., Dale, A. M. & Boas, D. A. Spatial extent of oxygen metabolism and hemodynamic changes during functional activation of the rat somatosensory cortex. Neuroimage 27, 279–290 (2005).

72. Fang, Q. & Boas, D. A. Monte Carlo Simulation of Photon Migration in 3D Turbid Media Accelerated by Graphics Processing Units. Opt. Express 17, 20178 (2009).

73. Chen, J., Fang, Q. & Intes, X. Mesh-based Monte Carlo method in time-domain widefield fluorescence molecular tomography. J. Biomed. Opt. 17, 1 (2012).

74. Sunil, S. et al. NeuroImage : Clinical Stroke core revealed by tissue scattering using spatial frequency domain imaging. NeuroImage Clin. 29, 102539 (2021).

75. Srinivasan, V. J. et al. Multiparametric, Longitudinal Optical Coherence Tomography Imaging Reveals Acute Injury and Chronic Recovery in Experimental Ischemic Stroke. PLoS One 8, (2013).

